# JAX Animal Behavior System (JABS): A genetics informed, end-to-end advanced behavioral phenotyping platform for the laboratory mouse

**DOI:** 10.1101/2022.01.13.476229

**Authors:** Anshul Choudhary, Brian Q. Geuther, Thomas J. Sproule, Glen Beane, Vivek Kohar, Jarek Trapszo, Vivek Kumar

**Affiliations:** The Jackson Laboratory, 600 Main Street, Bar Harbor ME 04609

**Author notes:** Contributed Equally (Anshul Choudhary, Brian Q. Geuther, Thomas J. Sproule, Glen Beane).

## Abstract

Automated detection of complex animal behavior remains a challenge in neuroscience. Developments in computer vision have greatly advanced automated behavior detection and allow high-throughput preclinical and mechanistic studies. An integrated hardware and software solution is necessary to facilitate the adoption of these advances in the field of behavioral neurogenetics, particularly for non-computational laboratories. We have published a series of papers using an open field arena to annotate complex behaviors such as grooming, posture, and gait as well as higher-level constructs such as biological age and pain. Here, we present our, integrated rodent phenotyping platform, JAX Animal Behavior System (JABS), to the community for data acquisition, machine learning-based behavior annotation and classification, classifier sharing, and genetic analysis. The JABS Data Acquisition Module (JABS-DA) enables uniform data collection with its combination of 3D hardware designs and software for real-time monitoring and video data collection. JABS-Active Learning Module (JABS-AL) allows behavior annotation, classifier training, and validation. We introduce a novel graph-based framework (*ethograph*) that enables efficient boutwise comparison of JABS-AL classifiers. JABS-Analysis and Integration Module (JABS-AI), a web application, facilitates users to deploy and share any classifier that has been trained on JABS, reducing the effort required for behavior annotation. It supports the inference and sharing of the trained JABS classifiers and downstream genetic analyses (heritability and genetic correlation) on three curated datasets spanning 168 mouse strains that we are publicly releasing alongside this study. This enables the use of genetics as a guide to proper behavior classifier selection. This open-source tool is an ecosystem that allows the neuroscience and genetics community for shared advanced behavior analysis and reduces the barrier to entry into this new field.

## 1 Introduction

Behavioral analysis in animal models seeks to link complex and dynamic behaviors with underlying genetic and neural circuit functions [1]. In the context of disease, altered genetic circuits shape altered neural circuits, which in turn produces altered behaviors. The primary purpose of behavior analysis in the animal is to understand the mechanisms of disease and to seek novel therapeutics to improve human health. The laboratory mouse has been at the forefront of these discoveries. However, linking altered genetic circuits to functional changes in neural circuits and ultimately behavior is challenging. These challenges are broad, however one major hurdle has always been a behavior quantification task itself. Animal behavior quantification has rapidly advanced in the past few years with the application of machine learning to the problem of behavior annotation and with the adoption of computational ethology approaches to behavioral neurogenetics [2–6].

These advances are mainly due to breakthroughs in the statistical learning and computer science fields which have been adopted and extended for biological applications, and have made the task of behavior annotation at high resolution scalable and objective, with increased accuracy. [7]. Although significant advances have been made in the annotation of animal behavior using machine vision, a major challenge remains in the democratization of these technologies. As a simple example, many labs adopt their existing apparatus to generate an intermediate representation of the animal for tasks such as tracking. These are often segmentation masks or keypoints. Each lab generally trains a custom model for these, which, depending on the complexity of the task, can require large amounts of human annotated training data. Many do not validate or even report the performance of their models, which are taken at face value to work. This is a large data labeling burden that is repeated by individual labs. The next step of extracting behaviors from these intermediate representations is even more challenging. The process entails creating features from intermediate representations followed by heuristics or classifiers to determine when a behavior of interest occurs. Behaviorists often disagree on behavior definition, even within labs, and therefore these behavior classifiers are incredibly valuable. They encode a behaviorist’s expertise in the form of mathematical weights. Since labs start with niche behavior apparatus and intermediate representations, the process of feature extraction, classification, the logic of assigning behaviors stays within a lab. That is, it is challenging for labs to share classifiers, because they only work in their hardware setup. This paradigm is not sustainable and prohibits the application of engineering principles to biology. The paradigm described above, combined with the fact that a high level of expertise is needed for proper use and interpretation of machine learning methods, can be a challenge to the reproducibility and replicability of scientific discoveries and ultimately therapeutic discoveries.

With this in mind, we present two complementary systems that are designed for behavior characterization in rodent models. The first platform, called JAX Animal Behavior System (JABS), consists of video collection hardware and software, a behavior labeling and active learning app, and an online database for sharing classifiers. This is an open field system which we have used in over 6 papers [8–13]. Adoption of JABS will allow laboratories to bypass the need for creating segmentation or pose estimation models for routine open-field tasks. In addition, existing models for frailty [14], nociception [13], seizures [15], and others can be adopted. The second, called Digital InVivo System (DIV Sys) is hardened and scalable home cage monitoring system (see Robertson et. al.). Both end-to-end systems are designed to enable community members to leverage others’ work and to extend the capabilities of the system. We hope that these platforms will be adopted and extended by the community.

JABS adds to a list of commercial and open source animal tracking platforms. JABS covers hardware, behavior prediction, a shared resource for classifiers, and genetic association studies. Were not aware of another system that encompasses all these components. Commercial packages such as EthoVision XT and HomeCage Scan give users a ready-made camera-plus-software solution that automatically tracks each mouse and reports simple measures such as distance traveled or time spent in preset zones, but they do not provide open hardware designs, editable behavior classifiers, or any genetics workflow. At the open-source end, there are greater than 100 projects cataloged on OpenBehavior and summarized in recent reviews (Luxem et al., 2023 [4]; Ik & Ünal 2023 [16]) that usually cover only one link in the chainDIY rigs, pose-tracking libraries (e.g., DeepLabCut [17], SLEAP [18]) or supervised and unsupervised behaviour-classifier pipelines (e.g., SimBA [19], MARS [20], JAABA [21], B-SOiD [22], DeepEthogram [23]). JABS provides an open source ecosystem that integrates all four: (i) top-down arena hardware with parts list and assembly guide; (ii) an active-learning GUI that produces shareable classifiers; (iii) a public web service that enables sharing of the trained classifier and applies any up-loaded classifier to a large and diverse strain survey; and (iv) built-in heritability, genetic-correlation and GWAS reporting.

## 2 Results

JABS is an integrated platform developed in our lab over the past five years with pose and segmentation models as intermediate representations. Our lab has previously used computer-vision methods to track visually diverse mice under different environmental conditions [8], infer pose for gait analysis and posture analysis [10], and detect complex behaviors like grooming [9]. We have also used computervision derived features to predict complex constructs, such as health and frailty and pain [11, 13]. These models have been trained and validated on genetically diverse mouse strains for high-quality foundational metrics [8, 10]. JABS hardware and software has been used to characterize complex behaviors such as grooming, gait, posture, as well as complex states such as frailty, pain, and intensity of seizures [8–11]. JAX has made components of JABS including ML models are free to use for non-commercial purposes.

The process and various components of JABS are illustrated Figure 1A. Briefly, our system comprises of three components encompassing five different processes, namely, i) data acquisition, ii) behavior annotation, iii) classifier training, iv) behavior characterization, and v) data integration. The first component (JABS-DA module) is the custom designed standardized data acquisition hardware and software that provides a controlled environment, optimized video storage, and live monitoring capabilities. The second component (JABS-AL module) is a python based GUI active learning app for behavior annotation and training classifiers using the annotated data. One can then use the trained classifiers for predicting whether behavior happens or not in the unlabeled frames. The last component of JABS is analysis and integration module (JABS-AI), a web application that provides an inter-active user interface to browse through the strain survey results from different classifiers, download existing classifiers and related training data. The app can also be used to classify various behaviors in user submitted videos (pose files) using the classifiers available in the database. Furthermore, researchers have the option to contribute their custom classifiers, trained through the JABS-AL app. These user-generated classifiers can be submitted to perform predictions within our extensive strain survey dataset, coupled with comprehensive genetic analysis, including assessments of heritability and genetic correlations. Next, we discuss the individual components of JABS in detail.

**Figure 1:**
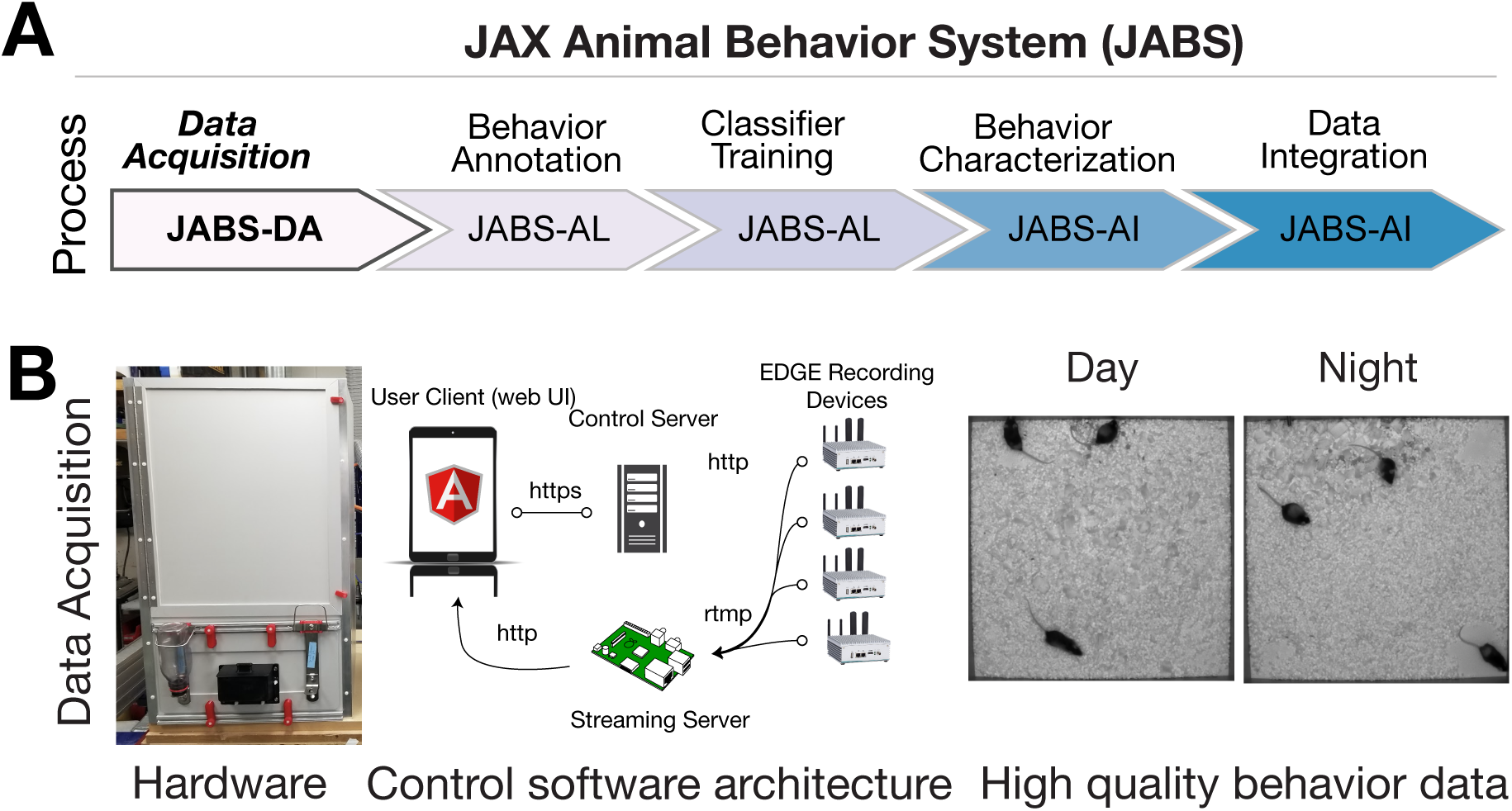
JABS data acquisition module (JABS-DA). The JABS-DA consists of hardware and software for video data acquisition and processing. (A) JABS pipeline highlighting individual steps towards automated behavioral quantification. (B) Detailed example of JABS data acquisition, including a picture of the monitoring hardware, architecture of the real-time monitoring app, and screenshots from videos taken during daytime and nighttime. The open field arena is shown from the outside (left), and a screenshot of video data is shown on the right. JABS-DA blocks visible light to the camera and only collects data using IR illumination, which produces uniform data during day and night. The JABS-DA computer hardware and software (middle) allow streaming of video data from edge devices, which enables remote welfare checks and web based experiment setup and monitoring. Data compression is handled on these edge devices.

### 2.1 JABS data acquisition - Hardware and Software

We use a standardized hardware setup for high quality data collection and optimized storage (Figure 1B). The end result is uniform video data across day and night. Complete details of the software and hardware, including 3D designs used for data collection, are available on our Github (https://github.com/KumarLabJax/JABS-data-pipeline/tree/main). We also provide a step-by-step assembly guide (https://github.com/KumarLabJax/JABS-data-pipeline/blob/main/Multi-day%20setup%20PowerPoint%20V3.pptx).

We have organized the animal habitat design into three groups of specifications. The first group of specifications are requirements necessary for enabling compatibility with our machine learning algorithms. The second group describes components that can be modified as long as they produce data that adheres to the first group. The third group describes components that do not affect compatibility with our machine learning algorithms. While we distinguish between abstract requirements in group 1 and specific hardware in group 2 that meet those requirements, we recommend that users of our algorithms use our specific hardware in group 2 to ensure compatibility.

The design elements that are critical to match specifications in order to re-use machine learning algorithms include (1a) the camera viewpoint, (1b) minimum camera resolution and frame rate, (1c) field of view and imaging distance, (1d) image quality and (1e) the general appearance of the habitat (cage or enclosure). The design elements that are flexible but impact the compatibility are (2a) camera model, (2b) compute interface for capturing frames, (2c) lighting conditions, (2d) strains and ages of mice and (2e) animal bedding contents and quantity. Design elements that have no impact on compatibility are (3a) height of habitat walls to prevent mice from escaping, (3b) animal husbandry concerns, (3c) mounting hardware, (3d) technician ergonomic considerations and (3e) electrical connection hardware and management.

#### 2.1.1 Group 1 specifications

Generally speaking, multi-view imaging will provide the most accurate pose models but requires increased resources on both hardware setup as well as processing of data. Our system operates on a top-down camera viewpoint. This specification enables flexibility and allows more diverse downstream hardware and ease of construction. Top-down provides the advantage of flexibility for materials, since the floor doesnt need to be transparent. Additionally lighting and potential reflection with the bottom-up perspective. Since the paws are not occluded from the bottom-up perspective, models should have improved paw keypoint precision allowing the model to observe more subtle behaviors. However, the appearance of the arena floor will change over time as the mice defecate and urinate. Care must be taken to clean the arena between recordings to ensure transparency is maintained. This doesnt impact top-down imaging that much but will occlude or distort from the bottom-up perspective. Additionally, the inclusion of bedding for longer recordings, which is required by IACUC, will essentially render bottom-up imaging useless because the bedding will completely obscure the mouse.

Our algorithms are trained using data originating from 800×800 pixel resolution image data and 30 frames per second temporal resolution. This resolution was selected to strike a balance between resolution of the data and size of data produced. While imaging at higher spatial and temporal resolution is possible and sometimes necessary for certain behaviors, these values were selected for general mouse behavior such as grooming, gait, posture, and social interactions. We train and test our developed algorithms against the spatial resolution. We note that these are minimum requirements, and down-sampling higher resolution and frame rate data still allows our algorithms to be applied.

Similar to the pixel resolution, we also specify the field of view and imaging distance for the acquired images in real-world coordinates. These are necessary to achieve similar camera perspectives on imaged mice. Cameras must be mounted at a working distance of approximately 100cm above the floor of the arena. Additionally, the field of view of the arena should allow for between 5 − 15% of the pixels to view the walls (field of view between 55cm and 60cm). Having the camera a far distance away from the arena floor reduces the effect of both perspective distortion and barrel distortion. We selected values such that our custom camera calibrations are not necessary, as any error introduced by these distortions are typically less than 1%.

Additionally, image quality is important for meeting valid criteria for enabling the use of machine learning algorithms. Carefully adjusting a variety of parameters of hardware and software values in order to achieve similar sharpness and overall quality of the image is important. While we cannot provide an exact number or metric to meet this quality, users of our algorithms should strive for equal or better quality that exists within our training data. One of the most overlooked aspect of image quality in behavioral recordings is image compression. We recommend against using typical software-default video compression algorithms and instead recommend using either defaults outlined in the software we use or recording uncompressed video data. Using software-defaults will introduce compression artifacts into the video and will affect algorithm performance. For example, most video recording software will default to having a constant keyframe rate, typically around every 30 frames. This will artificially create a measurable signal in video features at the keyframe rate.

Finally, the general appearance of the cage should be visually similar to the variety of training data used in training the machine learning algorithms. Documentation on this for each individual algorithm for assessing the limitations are published [8–11]. While our group strives for the most general visual diversities in mice behavioral assays, we still need to acknowledge that any machine learning algorithms should always be validated on new datasets that they are applied to. Generally our machine learning algorithms earlier in the entire processing pipeline, such as pose estimation, are trained on more diverse datasets than algorithms later in the pipeline, such as pain and frailty predictions.

#### 2.1.2 Group 2 specifications

In order to achieve compliant imaging data for use with our machine learning algorithms, we specify the hardware we use. While the hardware and software mentioned in this section is modifiable, we recommend that careful consideration is taken such that changes still produce complaint video data.

We modified a standard open field arena that has been used for high-throughput behavioral screens [24]. The animal environment floor is 52 cm square with 92 cm high walls to prevent animals escaping and to limit environmental effects. The floor was cut from a 6mm sheet of Celtec (Scranton, PA) Expanded PVC Sheet, Celtec 700, White, Satin / Smooth, Digital Print Gradesquare and the walls from 6mm thick Celtec Expanded PVC Sheet, Celtec 700, Gray, (6 mm x 48 in x 96 in), Satin / Smooth, Digital Print Grade. All non-moving seams were bonded with adhesive from the same manufacturer. We used a Basler (Highland, IL) acA1300-75gm camera with a Tamron (Commack, NY) 12VM412ASIR 1/2" 4-12mm F/1.2 Infrared Manual C-Mount Lens. Additionally, to control for lighting conditions, we mounted a Hoya (Wattana, Bangkok) IR-80 (800nm), 50.8mm Sq., 2.5mm Thick, Colored Glass Longpass Filter in front of the lens using a 3D printed mount. Our cameras are mounted 105 +/- 5 cm above the habitat floor and powered the camera using the power over ethernet (PoE) option with a TRENDnet (Torrance, CA) Gigabit Power Over Ethernet Plus Injector. For IR lighting, we used 6 10 inch segments of LED infrared light strips (LightingWill DC12V SMD5050 300LEDs IR InfraRed 940nm Tri-chip White PCB Flexible LED Strips 60LEDs 14.4W Per Meter) mounted on 16-inch plastic around the camera. We used 940nm LED after testing 850nm LED which produced a marked red hue. The light sections were coupled with the manufactured connectors and powered from an 120vac:12vdc power adapter.

For image capture, we connected the camera to an nVidia (Santa Clara, CA) Jetson AGX Xavier development kit embedded computer. To store the images, we connected a local four-terabyte (4TB) USB connected hard drive (Toshiba (Tokyo, Japan) Canvio Basics 4TB Portable External Hard Drive USB 3.0) to the embedded device. When writing compressed videos to disk, our software both applies optimized de-noising filters as well as selecting low compression settings for the codec. While most other systems rely on the codec for compression, we rely on applying more specific de-noising to remove unwanted information instead of risking visual artifacts in important areas of the image. We utilize the free ffmpeg library for handling this filtering and compression steps with the specific settings available in our shared C++ recording software. Complete parts list and assembly steps are described in (https://github.com/KumarLabJax/JABS-data-pipeline)

#### 2.1.3 Group 3 specifications

Finally, here we present hardware and software that can be modified without risk of affecting video compliance. For natural light, we used a F&V (Netherlands) fully dimmable R-300SE Daylight LED ring light powered by a 120vac:12vdc power adapter. These lights are adjustable to meet the visible lighting needs of specific assays without affecting the visual appearance of the data. To keep the animals nourished, we installed water bottles and a food hopper external to the animal environment. These were placed on the outside of the arena on a removable panel. The panel can be customized as needed for experiments without the need to replace/modify the entire arena. To suspend the camera and lights, we used a wire shelf from our solution for technician ergonomics.

To raise the animal cage to an ergonomic height, we used the 24-inch by 24-inch option of the Metro (Wilkes-Barre, PA) Super Erecta wire shelving system with three shelves. As mentioned in the earlier paragraph, the topmost shelf was used to suspend the camera and lights. We also hinged one wall, turning it into a door, to allow easier animal access. Communication between the electronic devices was interconnected with CAT5 cables and a network switch and a powered USB hub was used between the USB connected hard drive and the nVidia compute device. We used a digital timer for the visible LED light, a 120v power strip to consolidate the power, and a universal power source (battery backup) between the chamber and facility power.

For ease of use and reduction of environmental noise, we also include a software for remote monitoring and welfare check. The software consists of three main components: a recording client implemented in C++, a control server implemented with the Flask Python framework, and a web-based user interface implemented with Angular (Figure 1). The recording client runs locally on each Nvidia Jetson Xavier computer and communicates with the server using the Microsoft C++ REST SDK to provide centralized monitoring and control of distributed recording devices. The recording client captures raw frames from the camera and encodes video using the ffmpeg library. In addition to saving encoded video on the local hard drive, the recording client can optionally send video over the RTMP protocol to a NGINX server configured with the nginx-rtmp plug-in. The web interface communicates with the control server, which relays recording start and stop commands to individual recording devices, enabling the user to remotely control various aspects of recording in addition to viewing the live stream from the NGINX streaming server using the HTTP Live Streaming (HLS) protocol (Figure 2). Each JABS-DA unit has its own edge device (Nvidia Jetson). Each system (which we define as multiple JABS-DA areas associated with one lab/group) can have multiple recording devices (arenas). The system requires only 1 control portal (RPi computer) and can handle as many recording devices as needed (Nvidia computer w/ camera associated with each JABS-DA arena). To collect data, 1 additional computer is needed to visit the web control portal and initiate a recording session. Since this is a web portal, users can use any computer or a tablet. The recording devices are not strictly synchronized but can be controlled in a unified manner.

**Figure 2:**
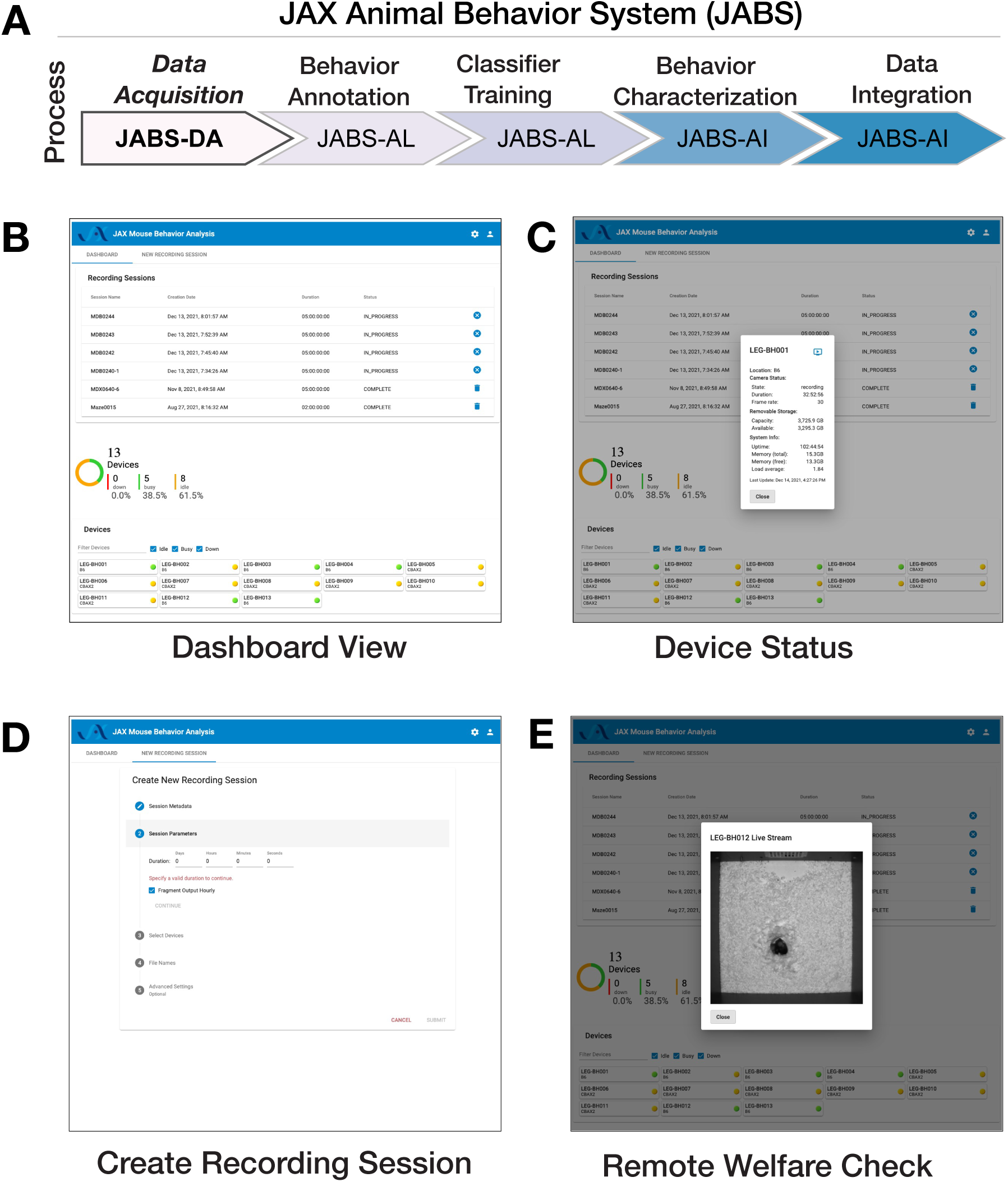
JABS data acquisition module (JABS-DA) consists of a web-based control system for recording and monitoring experiments. (A) JABS pipeline. (B-E) Screenshots from Angular web client that allows monitoring of multiple JABS Acquisition units in multiple physical locations can be seen on one screen (B). Dashboard view allows monitoring of all JABS units and their status, Device Status provided detailed data on individual devices (C) Recording session dashboard allows initiation of new experiments (D), and remote welfare view allows live video to be streamed from each unit (E).

### 2.2 Environment checks

To evaluate the suitability of JABS-DA for long-term housing of mice, we conducted a series of experiments comparing environmental conditions and animal health outcomes within these arenas to those observed in standard JAX housing cages. Our goal was to provide data for the JAX Institutional Animal Care and Use Committee (IACUC) to confirm health and welfare of animals over time in these apparatus. These data can be used for Institutional ACUC protocols by others. We compare our data with established guidelines from the Guide for the Care and Use of Laboratory Animals (the Guide) [25]. Our experiments were performed in one room at The Jackson Laboratory, Bar Harbor, Maine (JAX) with temperature and humidity set to 70-74*°*F (^∼^21-23*°*C) and 40-50%, respectively.

One concern related to use of the JABS arena in long-term experiments was that the 90 cm height of the walls without lower air openings might result in inadequate air flow and build up of toxic gases. To address this, we compared environmental parameters in JABS arenas with that of a standard JAX housing cage. Two JABS arenas were observed with 12 male C57BL/6J mice 12-16 weeks old in each for a 14-day period. At the same time, one wean cage containing 10 male C57BL/6J age-matched mice was observed on a conventional rack for matching air flow in the same room. We used a #2 Wean Cage (30.80 x 30.80 x 14.29 cm) from Thoren (Hazleton, Pennsylvania) with 727.8 cm^2^ floor space, which is a common housing container for mice and is approved at JAX to house 10 animals. This commercial cage has a floor area that is ^∼^1/4 that of the JABS arena. The ceiling height in the wean cage ranges 5-14 cm due to the sloped metal wire cover that contains food and water. The JABS arena, by contrast, has no ceiling. Food, water, bedding type and depth and light level were all matched in the arenas and wean cage. Bedding (1:1 ratio of aspen chip/shavings mix and Alpha-Dri) was left unchanged for the full two-week period to minimize interaction with mice in JABS arenas as much as possible. To determine if forced air flow was needed for an acceptable arena environment, one of the two arenas and the wean cage were exposed to normal room air flow, while the second arena had a 6-inch quiet electric fan mounted above for increased circulation. The fan was pointed to blow air up to draw air out of the arena instead of actively blowing air towards the mice.

We monitored CO_2_ and ammonia, common housing gases [25]. CO_2_ was measured with an Amprobe CO_2_ meter daily, excluding weekends and holidays, in both arenas and the wean cage. CO_2_ was recorded in the room’s center before and after each arena and wean cage measurement as a control. For higher levels, CO_2_ is shown as a range due to oscillation. Ammonia was tested with Sensidyne Ammonia Gas Detector Tubes (5-260 ppm) in the arena without a fan and the wean cage on days 0, 2, 4, 7, and 14, with samples taken near the floor and waste accumulation areas. Temperature and humidity data loggers (MadgeTech RHTEMP1000IS) were placed on the floor in each arena and the wean cage for the experiment’s duration. An environment monitor (Hobo, U12-012, Onset) was mounted on the wall for room background data. Body weight was measured daily, excluding weekends and holidays. Grain and water were weighed at the start and end of each experiment to check consumption.

We observed daily room background CO_2_ levels of 454 to 502 ppm throughout the 14-day experiment. These are very close to expected outdoors levels and indicative of a high air exchange rate [26]. JABS arena CO_2_ levels varied from a low of 511 ppm on day 1 to an oscillating range of 630 to 1565 ppm on day 14. The JAX standard wean cage experienced an oscillating range of 2320 to 2830 ppm on day 0 climbing to an oscillating range of 3650 to 4370 ppm on day 14. The wean cage CO_2_ values approximately match those from another published study of maximum grouped mice in similar static housing [27]. Indoor CO_2_ is often evaluated as level above background [26]. We observe a maximum JABS arena CO_2_ level above background of 1082 ppm. This is 3.8 fold lower than the maximum observed CO_2_ levels in the wean cage (4121 ppm) (Figure S1B, arena with fan excluded from graph for clarity).

Ammonia levels in the JABS arena were below 5 ppm on days 0, 2, 4, and 7, rising to 18 ppm on day 14. In the wean cage, levels were <5 ppm on days 0 and 2, rose to 52 ppm on day 4, and remained at ^∼^230ppm on days 7 and 14. Initial concerns about high JABS arena walls hindering airflow were alleviated as CO_2_ and ammonia levels indicated better air exchange than standard housing. NIOSH’s recommended maximum ammonia exposure for humans is 25 ppm over 8 hours, with a similar recommendation for mice [25, 28]. Ammonia levels are mainly influenced by air changes per hour (ACH) [29, 30]. JAX animal rooms have ^∼^10 ACH and PIV cages have ^∼^55-65 ACH. Ammonia levels were consistently 10-50 times lower in the JABS arena compared to the control static wean cage and remained well within recommended limits (Figure S1C). Future JABS arena observations must consider the impact of ammonia on behavior [31]. Mice used in JABS experiments come from PIV housing, where ammonia levels are expected to be similar to those in the JABS arena, minimizing behavioral impact [30].

Temperatures in all locations (room background, two JABS arenas and one wean cage) remained in a range of 22-26*°*C throughout the experiment. Variance in room background readings suggest temperature fluctuations are more due to innate room conditions (such as environmental controls) than anything else. We find that arena structure does not adversely affect control of the temperature to which mice are exposed (Figure S1D).

The probes which measured temperature also measured humidity. The room probe, mounted on a wall 1 foot above the floor in the 8×8 feet room, recorded consistent background humidity of 45 ±5% (Figure S1E, green line). Housing probes in the bedding of each chambercentered in JABS arenas and along a wall in the smaller wean cagerecorded 55-60% humidity in the JABS arenas, except for occasional spikes not correlated with background changes, likely due to mouse urination (Figure S1E, blue and black lines). In contrast, wean cage humidity rose from 55-60% to above 75% within 12 hours and continued climbing to 97.5% by day 14 (Figure S1E, red line). Higher humidity in the micro-environments was due to mouse urination and limited air flow (Guide [25]). The JABS arenas maintained a drier environment because they had a higher bedding to mouse ratio (3.2 times more per mouse) and better air circulation compared to the wean cage (Figure S1E).

Weight is often used as a marker for health though body condition score is used as a more reliable indicator of serious concerns [12, 32, 33]. Mice in JABS arenas lost weight compared to those in the wean cage and this was initially a cause of concern. However, mice in JABS arenas maintained a healthy appearance and normal body condition score throughout the experiment. Other measurements demonstrating normal parameters and other control experiments not shown additionally led us to believe the weight differences are because JABS arena mice are active while wean cage mice, with more limited movement available, are sedentary. Mice started the experiment at 25-33 grams body weight. The lowest average recorded during the experiment was 95.6% of the start value, for mice in the JABS arena without a fan on day 9. The lowest individual recorded was 85.8% of start value at 23.6 grams on day 14, also in the arena without a fan (Figure S1F).

Per mouse grain usage was comparable between the JABS arena and the wean cage and in an expected range [34] (Figure S1G). Per mouse water usage was comparable between the JABS arena and the wean cage and in the expected range [35]. Somewhat higher water use in the arena could be indicative of higher activity requiring more hydration (Figure S1H). Since only one JABS arena and one wean cage were tested, error bars are not available to aid in interpretation.

Three mice from one arena and three from a wean cage were necropsied immediately following 14 days in the JABS arena or control wean cage to determine if any environmental conditions, such as possible low air flow in arenas potentially leading to a buildup of heavy unhealthy gases like ammonia or CO_2_, were detrimental to mouse health. Nasal cavity, eyes, trachea, and lungs were collected from each mouse. They were H&E stained and analyzed by a qualified pathologist. No signs of pathology were observed in any of the tissue samples collected (Figure S2).

Based on these environmental and histological analysis, we conclude that the JABS arena is comparable and in many respects better than a standard wean cage. Lack of holes near the floor do not create a build up of ammonia or CO_2_. Mice ate and drank at normal levels. We initially observed a slight decrease in body weight, which is increased in the next few days. We hypothesize that this could be due to the novel environment and the increase in space for movement, leading to more active mice.

### 2.3 JABS-AL: An active learning module for behavior classifier training

In the section, we first present an overview on behavior annotation and classifier training using JABS-AL module which utilizes our python-based, open-source graphical user interface (GUI) application which has been developed to be compatible with Mac, Linux and Windows operating systems. We then evaluate the utility and accuracy of JABS trained classifiers through two complementary approaches. In the first approach, we benchmark the performance of JABS classifiers against a previous neural network based approach [9], providing us a comparison of the performance of the two approaches on the same dataset. In the second approach, we studied how classifiers for the same behavior trained by two different human annotators in the lab compare with each other in terms of behavior identification, allowing us to assess the inherent variability among expert annotators.

#### 2.3.1 Behavior annotation and classifier training

There are two prominent approaches in the literature for training behavioral classifiers. The first approach trains the classifiers using the raw video files, as previously demonstrated to identify grooming behavior through the use of a deep neural network [9, 23]. The second approach involves first extracting pose keypoints in each frame using deep neural networks, which serves as inputs for machine learning classifiers [17, 18, 36]. Previously, we utilized a deep neural network based classifier to extract poses and used the keypoints to study gait behavior [10]. Pose based approach offers the flexibility to use the identified poses for training classifiers for multiple behaviors and we used this approach for JABS. Additionally, the extracted keypoints can also be used to generate quantifiable and interpretable features that can be used to study various aspects of animal behavior such as gait and posture. In addition to the raw video file, JABS annotation and classification active learning module requires pose files from our previously established neural network for pose estimation as an input to train the classifiers. Note that the raw videos are needed only for annotating behaviors, and one can predict the behaviors using only the pose files.

We have developed an easy to use open source python GUI software to annotate behaviors in videos, as shown in Fig. 3A. This tool allows users to easily annotate behaviors in video recordings through mouse/trackpad or keyboard shortcuts, as well as the option to leave frames unlabeled for ambigious cases. The GUI provides statistics of the total number of frames as well as the number of frames and bouts annotated for a particular behavior. The annotations are displayed below the video as an ethogram (Fig. 3B).The user can annotate multiple behaviors for the same video. Once minimum number of frames (100) and videos (2) have been annotated, the user can train a classifier using either of the tree-based methods such as Random Forest (RF)/Gradient Boost/XGBoost (XGB) [37–40] and check the classifiers accuracy with k-fold cross-validation, selecting a value of k that balances computational efficiency and accuracy. The input features for these classifiers are derived from the animal’s pose, which must be estimated prior to using the GUI. This is accomplished using our separate pose tracking pipeline (https://github.com/KumarLabJax/mouse-tracking-runtime), which employs an HRNet-based neural network [10] to identify the location of twelve keypoints in each video frame. This pipeline is described in greater detail in the Methods section. We then compute a number of informative features like distance between various keypoints, linear and angular velocity between keypoints, etc. that are used as input for these classifiers. We also incorporate temporal information from the videos by computing window features that include information from *w* (window size) frames on each side of the current frame. A complete list of base features currently included in JABS is provided in the supplementary information (Table S4). The weights of different features used by the trained classifiers improve the interpretability of the classifiers.

**Figure 3:**
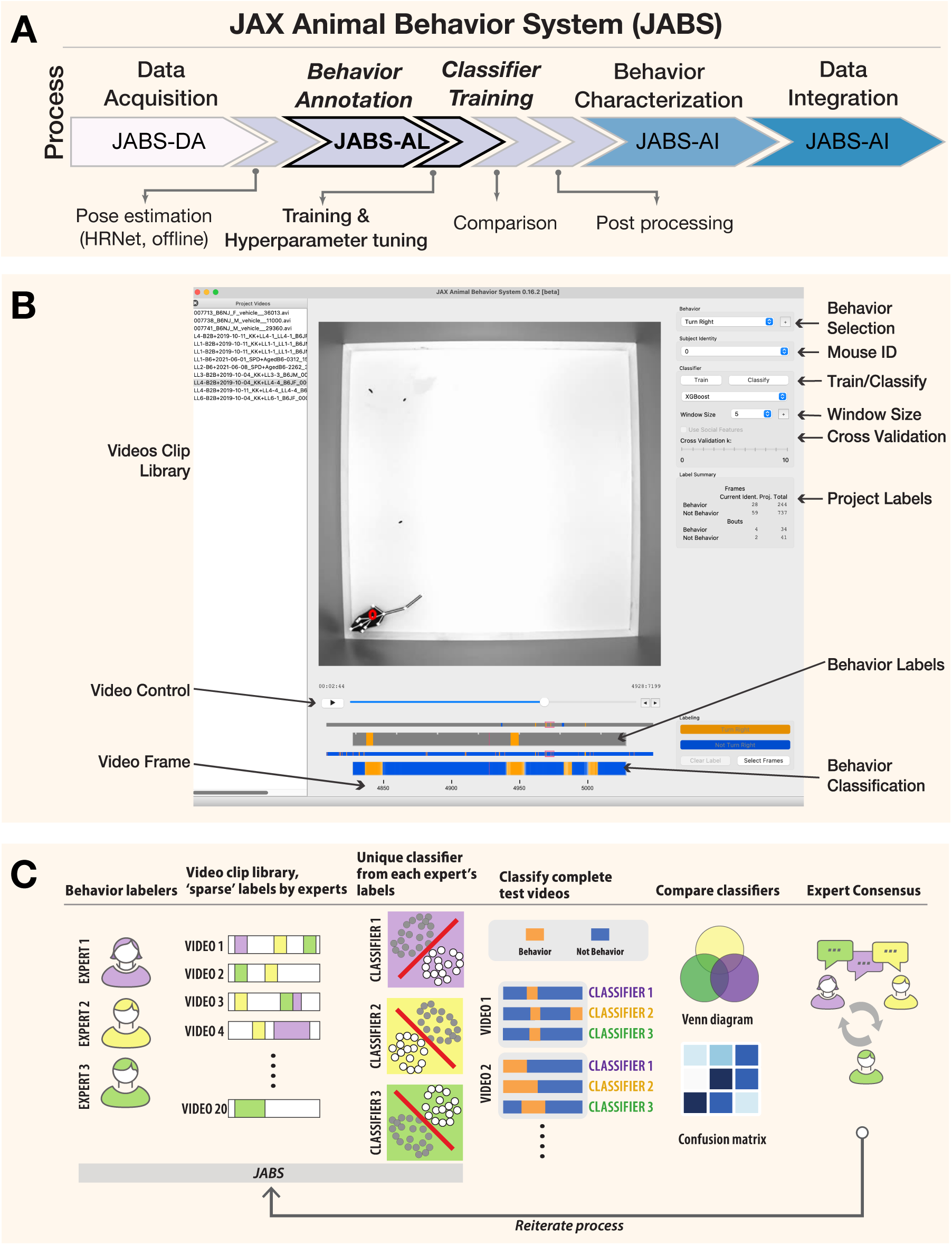
JABS-AL is a behavior annotation and classification module that allows training classifiers with sparse labels. (A) JABS pipeline highlighting individual steps towards automated behavioral quantification. (B) Screenshot of the python based open source GUI application used for annotating multiple videos frame by frame. One can annotate multiple mouse and for multiple behaviors. The labeled data is used for training classifiers using either random forest or gradient boosting methods. Adjustable window size (number of frames on the left and right of the current frame) to include features from a window of frames around the current frame. The labels and predicted labels are displayed at the bottom. (C) A sample workflow for training a typical classifier. Multiple experts can sparsely label videos to train multiple classifiers for the same behavior. These classifiers can be compared and experts can consult to iterate through the training process

Typically, to arrive at an optimal classifier for a behavior, we start by training multiple classifiers using annotated data from different human experts for the same set of videos and then evaluating performance of each classifier against a separate set of test videos as depicted in Fig. 3C. However, a core challenge is that different researchers may "see" and define the same behavior differently. This leads to low inter-rater reliability, where two people annotating the same video produce different labels. Disagreements often arise from two primary sources: the granularity of the labels and their reliance on subjective interpretation versus objective, physical descriptions. These are often referred to as labeling style due to annotator encoding their intuition about the behavior [20, 21]. Even when there is high interannotator agreement, explainable machine learning methods have shown that labelers rely on different features for labeling the same behavior [19]. Added frustration results when experts have to redefine and relabel videos to increase interannotator agreement. Researchers often try to break down complex behaviors into heuristic rules for the labelers to follow, which almost always leads to conflict between these rules and intuitive understanding of the complex behavior. These predetermined rules never capture the full repertoire of a complex behavior and often ask the labeler to violate their intuition.

In JABS we advocate that each expert sparsely labels frames from a video library to create a behavior classifier. These are essentially expert specific classifiers that encode the intuition or labeling style of that behaviorist. These classifiers are inferred on a set of test videos that have been set aside (Fig. 3C). Each expert’s classifiers are inferred from these test videos. We can then compare interannotator agreement using these inferred sense labeled videos. Experts can confer among themselves to reach a consensus on whether they agree on the inference of each classifier, potentially altering their definition of the behavior and altering their labeling style (Fig. 3C). JABS allows the experts to reiterate through the labeling process to arrive at a consensus. The process gives expert labelers an opportunity to agree on a behavior definition but does not operationalize the behavior through rules. In the end there may be multiple expert-specific classifiers for the same behavior, and end users must decide which classifiers they prefer to use. In this paper, we propose the use of broad-sense heritability to prioritize classifiers for the same behavior derived from different expert’s labels.

After classifiers are finalized, we often ask experts to densely label (label every frame) a another small set of test videos, which are used for final validation. We find that creating the dense label dataset at the end allows the labelers to have a clear understanding and intuitive definition of the behavior, whereas creating the dense label in the beginning of a project leads to frequent shift in behavior definition as the expert sees more instances of a behavior, particularly edge cages. The detailed user guide, along with a video tutorial on how to install and run the JABS Active Learning app, is available online (https://jabs-tutorial.readthedocs.io/en/latest/JABS_user_guide.html).

#### 2.3.2 Benchmarking JABS classifier using grooming behavior

Previously, a CNN based grooming behavior classifier trained on raw videos attained human level accuracy [9]. We re-purpose this large training dataset as a benchmark for estimating learning capacity of pose-based classifiers. For context, we report JABS classifier performance against both JAABA and CNN in [9]. Further, we evaluate how the performance of the classifier varies with the choice of machine learning algorithm, window size (*w*) of the features and the amount of training data. For the choice of machine learning algorithm, we utilize two popular tree-based methods, namely Random Forest (RF) and XGBoost (XGB). Briefly, the dataset contains 1,253 video segments, and we held out 153 video clips for validation (this is the same validation set used in [9]) and the rest are used to train the classifier. This split results in similar distributions of frame-level classifications between training and validation sets. More details of the dataset are available in Table-S1. We trained multiple classifiers by varying the amount of annotated data, window size, and machine learning algorithm. Our best accuracy from the neural network based approach for this dataset was 0.937 and the best classifier from JABS using all the annotated data, a window size (*w*) of 60 frames, and XGB machine learning algorithm achieved a comparable per-frame accuracy score of 0.9364. We noticed that with the same set of features, XGB typically achieved better accuracy than RF method across different window sizes and training data size. The results for these benchmark tests are shown in Figure 4B-D. Our tests with different window sizes show that grooming performance increases as we increase the window size, reaches a maximum (around 60 frames) and then degrades for large window sizes (Fig. 4B). Because grooming typically lasts for few seconds, classifiers using features within nearby frames will perform better as they incorporate optimal temporal information and including features from too few or too many frames will decrease the performance. We also investigate the impact of the amount of labeled data on the performance of JABS classifiers, as it can help to optimize the annotation process, ultimately reducing the time and resources required to train the model. To do this, we trained the XGB and RF classifiers using a subset of the full dataset (about 20 hours) consisting of 10, 20, 50, 100, 500 and 1100 training videos. These correspond to approximately 1.3%, 2.2%, 4.4%, 8.5%, 46.1% and 100% out of a total of 2181790 frames. As expected, the performance of JABS improves as we include more labeled data. However, the results demonstrate that a high degree of accuracy, approaching 85%, can be attained through the utilization of only 10 videos of training data, as evidenced by the corresponding area under the receiver operating characteristic curve (AUROC) of approximately 0.94, as depicted in Figure 4C-E. Additionally, it was found that the true positive rate (TPR) experienced a minimal decrease of about 1% when the training data was reduced from 100% to 50%, while maintaining a false positive rate (FPR) of 5% (Fig. 4F). To assess the generalizability of our grooming classifier across genetic diversity, we trained the model on videos spanning 60 genetically diverse mouse strains (n = 1,100 videos) and evaluated performance on an independent test set comprising 51 genetically diverse strains (n = 153 videos). Per-strain analysis revealed robust and uniform performance across the majority of genetic backgrounds (median F1 = 0.94, IQR = 0.8990.956). We observed only modest performance declines in albino strains, attributed to reduced visual contrast under infrared illumination conditions.

**Figure 4:**
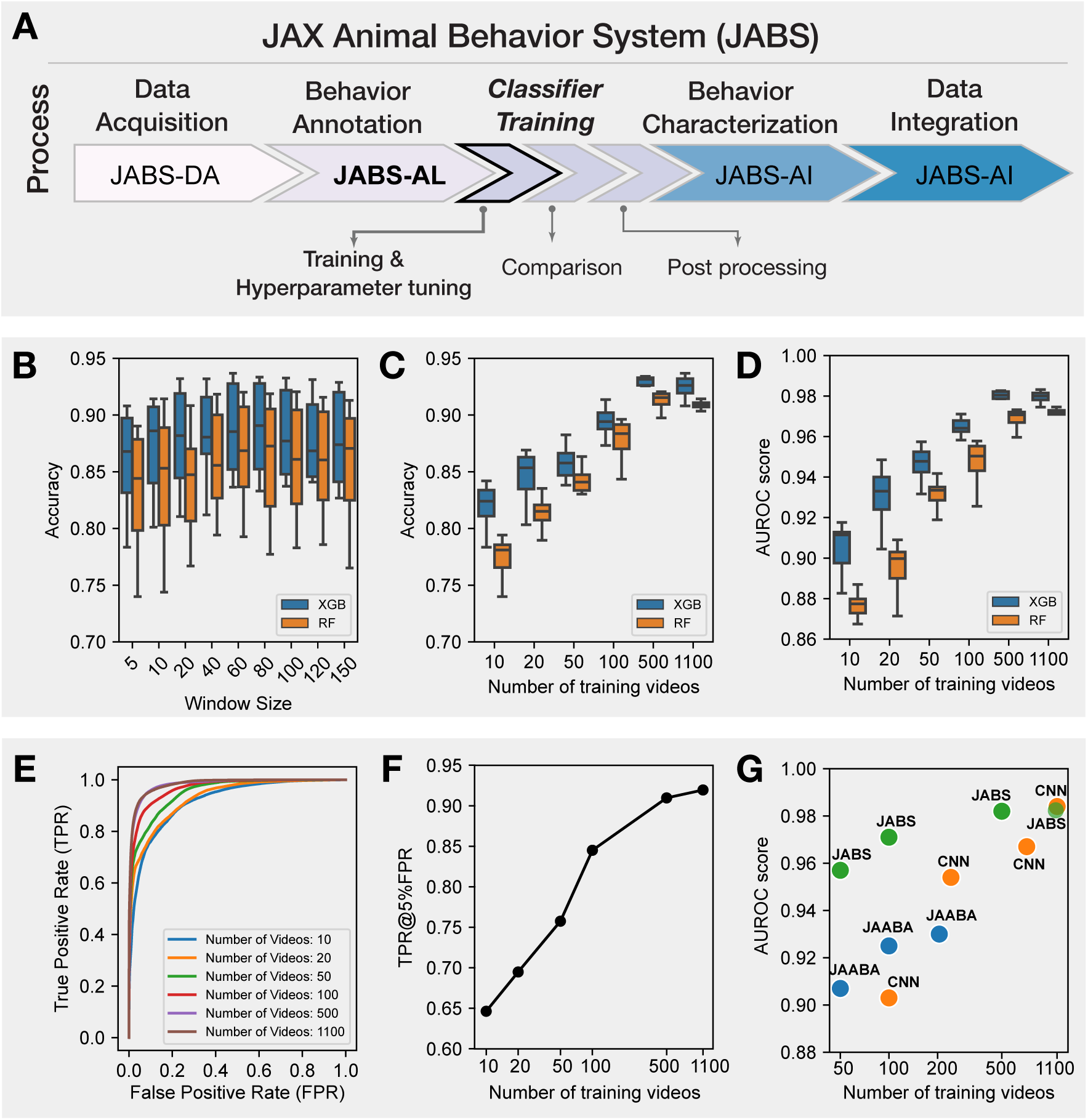
JABS Benchmarks: Selecting hyper-parameters and benchmarking JABS classifiers using grooming dataset. (A) JABS pipeline highlighting individual steps towards automated behavioral quantification. Using feature window size, type of classification algorithm and the number of training videos as our benchmarking parameters: (B) Accuracy of JABS classifiers trained using different window size (in frames) features. Each boxplot shows the range of accuracy values for different number of training videos and type of classification algorithms. (C, D) The effect of increasing the training data size on Accuracy and AUROC score of the JABS classifiers. (E) ROC curves for the JABS classifier trained with the window size of 60, XGB algorithm and varying training data size. (F) True positive rate at 5% false positive rate corresponding to the JABS classifier from panel (E) as the amount of training data is changed. (G) Comparing the performance of JABS based classifiers with a 3D Convolutional neural network (CNN) and JAABA based classifiers for different training data sizes. JAABA and CNN results were adopted from [9].

In the rapidly evolving field of automated quantification of animal behavior, two predominant methodologies have been established for learning behavior: using raw video data and using a reduced representation (abstraction) of the animal with certain keypoints, from which informed features are calculated [10, 19–21, 41]. To understand the trade-offs and strengths of each approach, we evaluate the performance of different classifiers that employ these methodologies when utilizing varying amounts of training data, as depicted in Fig. 4G. Interestingly, our findings demonstrate that utilizing keypoint-based low dimensional representation of animal behavior, as employed by JABS and JAABA [21] methodologies, leads to superior performance when compared to using high dimensional raw video data as employed by 3D CNNs, particularly when the availability of training data is limited. However, as the quantity of training data increases, the performance of both approaches tend to converge.

Therefore, by distilling the essence of a video into a series of key poses, JABS is able to effectively learn and generalize, even with smaller training sets. It has been shown to have a learning capacity on par with deep neural networks, as demonstrated by per-frame accuracy using the same benchmark data-set. Further, achieving 85% accuracy with just 1.4% of the labeled data, suggests that researchers can strike a balance between labeling efforts and desired accuracy by carefully selecting the amount of training data.

### 2.4 JABS analysis and integration module

In supervised machine learning, the accuracy and reliability of a trained classifier depends heavily on the quality of labeled data. Further, it has been observed that labeling of the same behavior by different human experts introduces variability among annotations due to variety of factors, including personal biases, subjectivity, and individual differences in understanding what constitutes a behavior [42, 43]. Therefore, it is critical to accurately capture the inter-annotator variability before selecting classifiers for downstream predictions. To capture this variability, we employ both frame based and bout based comparison and demonstrate that bout-based comparison gives a better estimate of inter-annotator agreement.

#### 2.4.1 Frame and bout-wise classifier comparison of inter-annotator variability

In order to test inter-annotator variability, we generated a set of single mouse behavior classifiers for two simple behaviors, left and right turn. We inferred behavior from all four classifies on a large set of videos and compared the two pairs of classifiers from each annotator (Figures 5, 6). The classifiers for all behaviors achieved good accuracy and F1 scores (Table S5). Further, the classifiers for the same behavior trained with different human annotations resulted in inter-annotator variability in predictions. This inter-annotator variability can be associated with (a) subjective differences of behavior definition among human labelers (b) varying level of annotator’s expertise, and (c) training with-in and across labs. We investigated the source of this variability and sought to determine the best method to mediate its effects. To capture this effect, we first visualized the predictions made by two classifiers trained for the same behaviors (left and right turn) but with different human annotators: annotator-1 (A1) & annotator-2 (A2). Figure 5B,C shows two sample ethograms corresponding to the predictions made by A1 & A2 for the left turn behavior. These ethograms show high level of concordance between the two annotators. However, upon closer examination, we observed that the percentage of left or right turn behavior predicted (for all the videos) by A2 was higher than A1 (see Figure 5D,G). The confusion matrix (shown in Figure 5E,H) quantifies the level of agreement between predictions made by annotators A1 and A2 for left and right turn behavior. However, since this behavioral task is heavily class-imbalanced (the number of frames with no-behavior is much more than that of behavior), accuracy can be misleading, as the classifier can achieve high accuracy by simply predicting majority class (not behavior) for all the frames. To address this imbalance, we calculate Cohen’s kappa (κ) metric [44] which is a commonly used measure of inter-annotator agreement accounting for the class imbalance. Mathematically, it is defined as 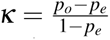 where *p_o_* is the observed agreement between annotators and *p_e_* is the expected agreement due to random chance. A κ score of 0 indicates that the agreement is no better than chance, and a score of 1 indicates perfect agreement, regardless of high/low accuracy. Finally, we visualize the frame-wise comparison of the two annotators showcasing the percentage of frames where the annotators agree and disagree on the occurence of a behavior as shown in Figure 5F,I. The venn diagram clearly highlights the discrepancy between high accuracy resulting from class imbalance (Figure 5E,H) and significant mismatch between % of predicted behavior (Figure 5D, G), with annotator A2 account for majority of discrepancy by predicting more frames as turning behavior compared to annotator A1.

**Figure 5:**
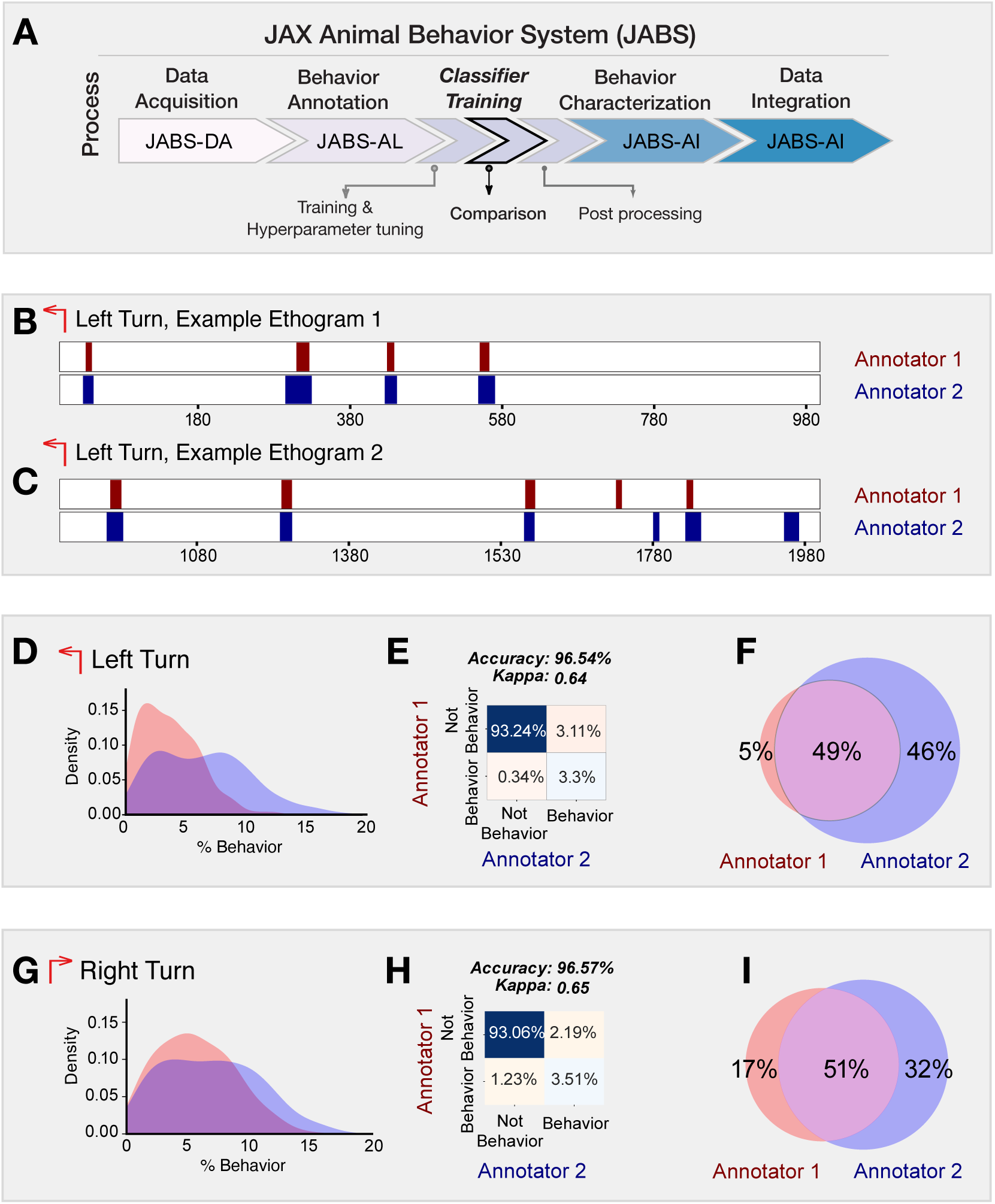
**Frame based comparison of classifiers from different annotators but trained for the same behavior**. (A) JABS pipeline highlighting individual steps towards automated behavioral quantification. (B, C) Two sample ethograms for the left turn behavior showing variation in behavior inference for two different annotators. (D, G) Kernel density estimate (KDE) of the percentage of frames predicted to be a left turn and a right turn respectively, by each annotator across all the videos. The major discrepancy between the two annotators is that A-2 systematically predicts larger number of frames as behavior compared to A-1. (E, H) Confusion matrix showing the agreement between predictions of two classifiers over all the videos in the strain survey for left and right turn behavior. (F, I) Venn diagram capturing the frame-wise behavior agreement between the two annotators for left and right turn behavior.

We observed in the ethogram (Figure 5B,C) that although many of the same bouts are captured by both A1 and A2, most of the frame discrepancies seem to be in the beginnings and ends of the bout. A2 seems to predict longer bouts than A1 (Figure 5D). Between two humans labeling the same behavior, there are unavoidable and sometimes substantial discrepancies in the exact frames of the behavior labeled even when trained in the same lab [20, 43]. To most behaviorists, detecting the same bouts of behavior is more important than the exact starting and ending frame of these bouts– as again, there are human-level discrepancies in this as well. Therefore, we used a bout-based comparison rather than a frame-based comparison to evaluate the performance of the classifiers.

For the bout-based comparison, we looked at how much overlap there was between the bouts of a behavior predicted by annotators A1 and A2, taking inspiration from the machine learning image-recognition and action-detection fields, where an overlap of pixels of the bounding box and ground truth label box called the intersection over union (IoU) [45, 46]. We developed a graph-based approach called an *ethograph* to represent the bouts of behavior recorded in the ethograms of annotators A1 and A2. Concretely, we define the ethograph for two annotators as a bipartite graph *G* = (*U,V, E*), where *U*, *V* are two disjoint sets corresponding to bouts predicted by each annotator and *E* represents the edges that connect each element in set *U* to an element in set *V* capturing the overlap in time between the bouts. Further, the vertices of an ethograph represents bouts with vertex color encoding for the annotator and vertex size proportional to the duration of the bout. Further, the edges (*E*) of the graph (*G*) represents the temporal overlap between the bouts (corresponding to different annotators) with the thickness of the edge proportional to the amount of bout overlap. Figure 6B,C shows the ethograms and their associated ethograph for the left turn behavior as predicted by annotators A1 and A2. In contrast to traditional frame-based ethograms, which simply display the sequential list of frames in which a behavior is observed, the ethograph allows for a more intuitive and visual representation of the temporal overlap between the bouts corresponding to different annotators (or even behaviors). This can be especially useful in identifying patterns and trends that may not be immediately apparent from comparing ethograms. By coloring the vertices and edges based on the annotator, it becomes easy to see which behaviors are consistently identified by both the annotators and which are more subjective and open to interpretation. Moreover, we can easily compute the bout-based agreement between the two annotators as the fraction of edges having thickness greater than some fixed threshold (see figure 6F for mathematical definition) which essentially means the fraction of bouts having overlap greater than a chosen overlap threshold. The bout agreement between two annotators for the left and right turn at a threshold of 0.5 is shown as a Venn diagram in Figure 6D,E along with the density distribution of bout length. The agreement between two annotators with bout-based measure was certainly much better than that with frame-based comparison (see figure 5F,I).

**Figure 6:**
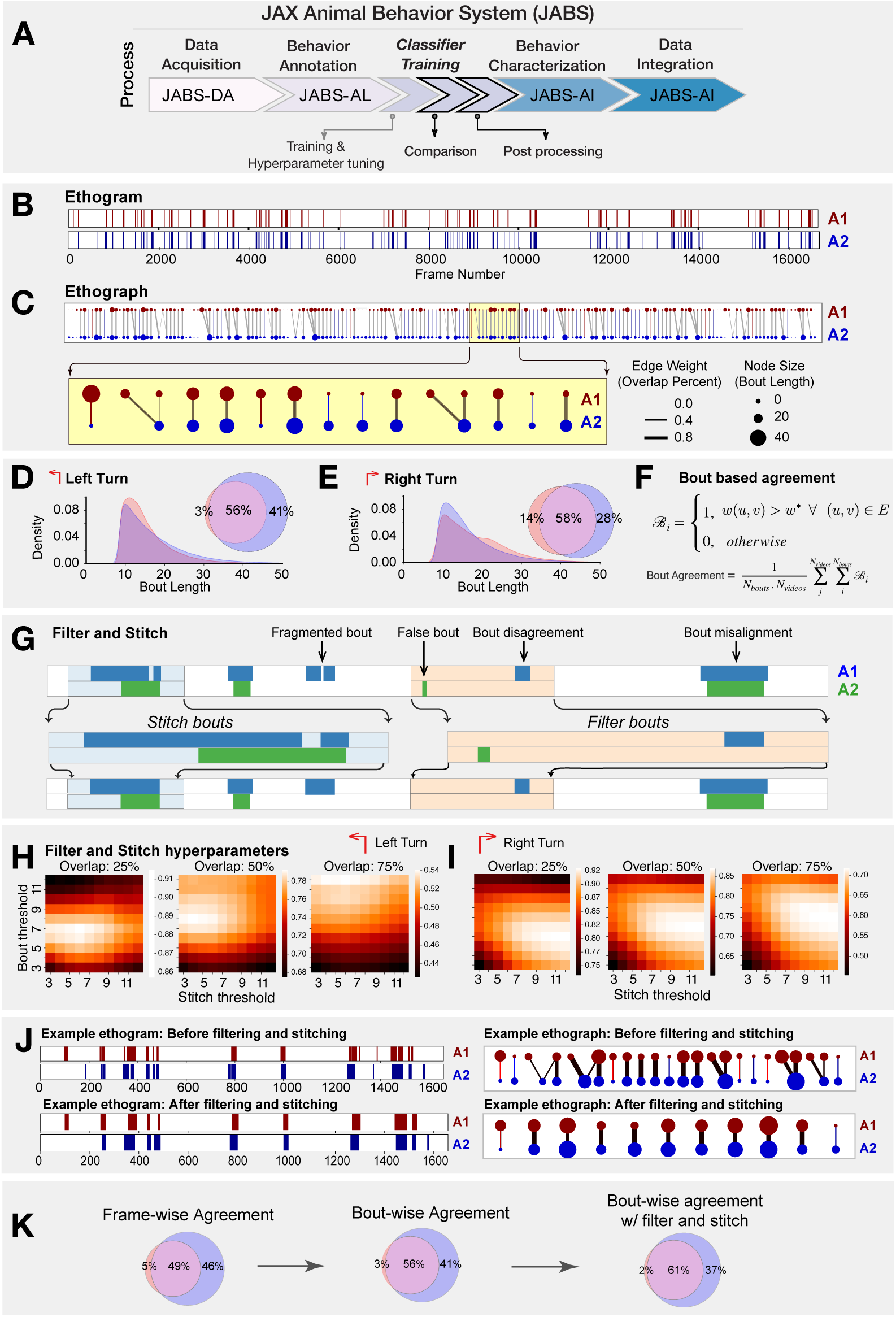
**Bout based comparison of classifier predictions from different annotators but trained for the same behavior**. (A) JABS pipeline highlighting individual steps towards automated behavioral quantification. (B) Ethogram depicting frame-wise left turn predictions for annotators A1 (red) and A2 (blue). (C) Ethograph corresponding to the ethogram in panel (B) capturing the bout level information as a bipartite network. The nodes represent bouts with node size & color proportional to the bout length & annotator respectively. Edge weights captures the fraction of bout overlap between two bouts predicted by different annotators for the same behavior. Edge weight and node size with zero value indicate missed bouts by an annotator. These have been given a small positive value for visualization purposes only. (D-E) Bout length distribution of annotators A1 & A2 for left and right turn behavior. (F) The mathematical definition of the average bout agreement between two annotators, where *w*(*u, v*) represents weight between nodes *u* and *v* (*u* ⊂ *U*, *v* ⊂ *V*) in the ethograph *G* (*U,V, E*) and *w*^∗^ is the bout overlap threshold (0.5 fixed for our study). (G) overview of the workflow for stitching and filtering at the bout level. (H, I) Hyper-parameters tuning to find optimal filtering and stitching thresholds. (J) Sample ethogram and its corresponding ethograph before and after applying stitching and filtering. (K) Inter-annotator agreement in frame wise predictions underestimates the agreement whereas the bout wise comparison post filtering and stitching captures the overall agreement in a more biologically meaningful way.

The predictions coming out of a classifier contained many short bouts (1-3 frames) of behavior that signal false positive bouts as they are much shorter than a typical bout of annotated behavior. Moreover, certain bouts of behavior were split by very short bouts (1-3 frames) of not-behavior signalling the presence of false negative bouts that results in fragmentation of a bout of behavior (see figure 6). To address this issue, we proposed a stitching and filtering step on the predictions coming out of classifier. First, we stitched those bouts whose distance to the neighboring bout is less than certain fixed threshold. This stitched the fragmented bouts as illustrated in Figure 6G. We then applied bout filtering which removed bouts of a length below a fixed threshold. To decide the optimal values of stitching and bout filtering thresholds, a hyper-parameter scan was performed for each behavior. Figure 6H,I presents the results from hyper-parameter scan over stitching and bout filtering thresholds when the value of percentage bout overlap is fixed at 25%, 50% and 75% for left (H) and right turn (I). Figure 6J captures the effect of applying bout filtering and stitching to a portion of an ethogram corresponding to the predictions made by A1 & A2 for the left turn behavior. The effect was clearly discernible when looking at the changes in ethograph, particularly with bouts (nodes) having multiple overlaps (edges) reducing to single overlap (edge) per bout.

In summary, when comparing classifiers, it’s important to consider the inherent variability of human annotators. Frame-wise comparison penalizes this natural variability, making it a sub-optimal measure of agreement. On the other hand, bout-wise comparison takes this variability into account, making it a more biologically meaningful measure of agreement between classifiers. In addition to using bout-wise comparison, applying techniques like stitching and filtering can further improve agreement by reducing false and fragmented bouts in classifier predictions. By considering these factors, we can better understand the inter-annotator variability and design more effective guidelines for behavior annotation.

#### 2.4.2 Compilation of Strain Survey Datasets

In the present study, we have curated and are releasing three comprehensive datasets to the public, namely JABS600, JABS1200 and JABS-BxD, that encapsulates behavioral data derived from approximately 168 unique mouse strains, ensuring a balanced representation with a nearly equable distribution between female and male sex. The JABS600 dataset includes a total of 598 videos corresponding to 60 strains, approximately balanced with five males and five females per strain. On the other hand, the JABS1200 dataset contains 1139 videos corresponding to 60 strains, representing approximately nine males and nine females represented per strain. Finally, the JABS-BxD dataset includes a total of 1083 videos corresponding to 108 BxD strains that are derived from a cross between C57BL/6J mice (B6) and DBA/2J mice (D2). The duration of each video is approximately one hour, furnishing a substantial repository of behavioral data, which is invaluable for large-scale automated analysis of behavioral patterns. Furthermore, each video is supplemented with a corresponding keypoint file comprising of 12 keypoints per frame, which is instrumental in extracting specific behavioral features. Additional information about the dataset is given in supplemental figure 2. In line with our dedication to scientific openness and collaboration, we have made these datasets - encompassing both the video recordings and the keypoint files - available for public access(https://dataverse.harvard.edu/ dataset.xhtml?persistentId=doi%3A10.7910%2FDVN%2FSAPNJG), making it easier for fellow researchers across labs to leverage our findings, replicate our experiments, and advance the field of automated behavior quantification.

#### 2.4.3 Strain Survey of Multiple Behaviors

One of the advantages of a standardized data acquisition system such as JABS is that data can be repurposed. For instance, a classifier trained by another lab could be inferred on videos generated by another lab. We trained a set of behavior classifiers using JABS active learning system and then inferred them on a previously published strain survey dataset [8]. The training dataset was composed of multiple human-annotated short videos (around 10 minutes each), we trained classifiers for left turn, right turn, grooming, rearing supported, rearing unsupported, scratch and escape as examples. These can easily be extended to other behaviors. To capture the effect of genotype on the behavior, we subsampled the original strain survey data set to 600 one-hour open field videos representing 60 different strains with 5 female and 5 male for each strain and make predictions using the trained classifiers. Further, we define 3 aggregate phenotype associated with each behavior namely the total duration of the behavior (in minutes) for the first 5, 20 and 55 minutes of the one-hour video [9], to capture the dynamic changes in behavior over time. The results are shown in Figure 7B, where the heatmap shows the Z-scores for the total duration of the behavior in 5, 20 and 55 minutes (|Z-score| *>* 1 thresholding is applied for easier visualization). The red and blue colored entries for a particular phenotype represents strains exhibiting the behavior that is more than one standard deviation above and below the mean of the phenotype respectively. Such data can have multiple utility. First, any user of JABS can conduct a rich analysis with little effort to yield biological insight. Such data can be used to refine classifiers by adding edge cases to training data. In addition, downstream genetic analysis suchs as heritability quantification and GWAS analysis are possible with this data [9, 10]. In our analysis, we observed a high number of escape attempts in C58/J mice. This strain has been shown previously to have high number of repetitive behaviors, perhaps even a strain for the study of autism features [47, 48] (Figure 7 Bottom panel). We find that other strains such as I/LnJ, C57/L, and MOLF/EiJ show increased levels of escape behaviors, thus increasing potential strains that could be used to model this behavior.

**Figure 7:**
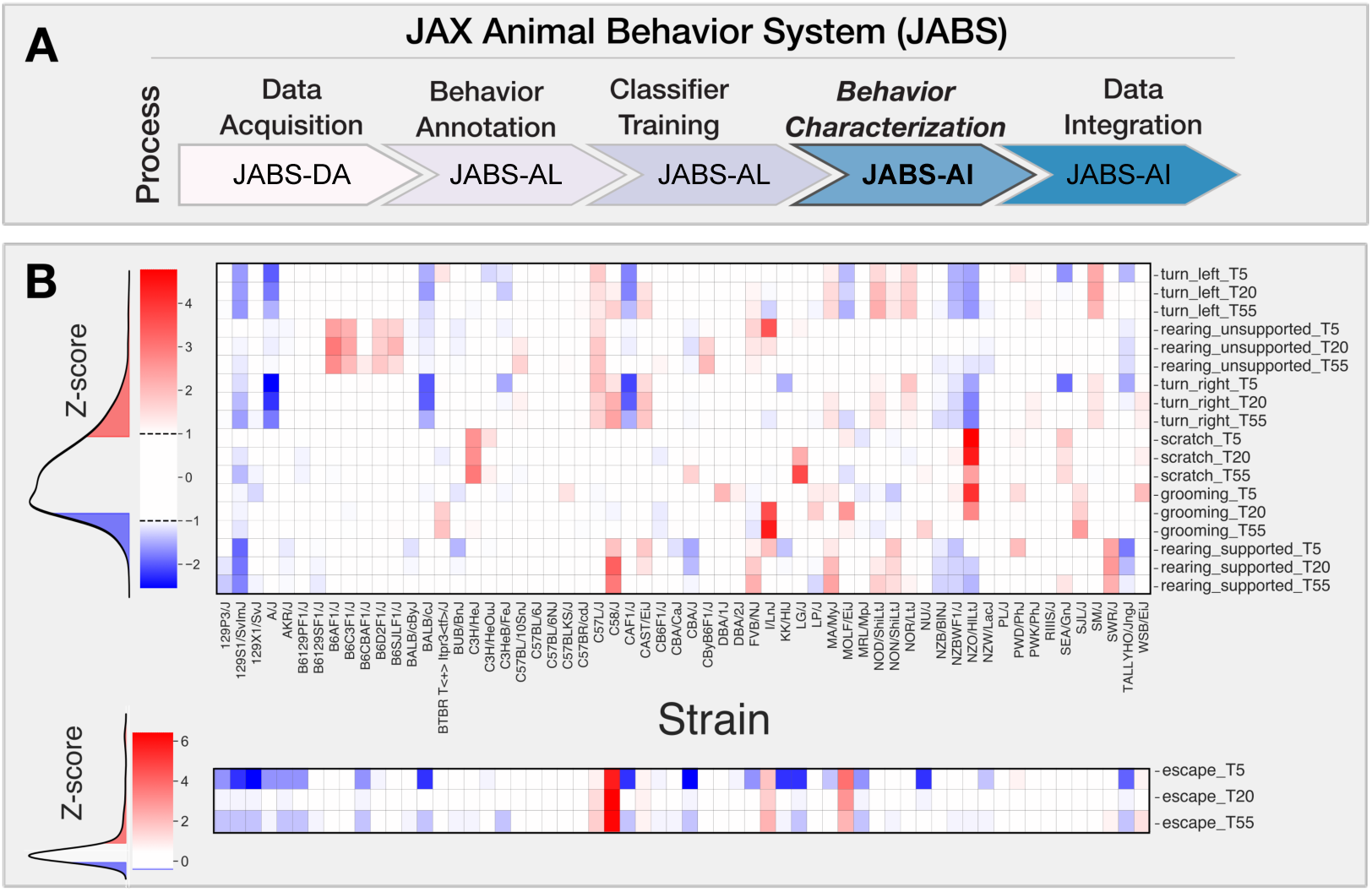
**JABS-AI (Analysis and Integration) module**: Strain-level behavioral phenotyping across genetically diverse mouse populations. (A) JABS pipeline highlighting individual steps towards automated behavioral quantification. (B) Heatmap showing Z-transformed behavioral scores for aggregate phenotypes measured at three time points (5, 20, and 55 minutes) across the JABS600 strain survey. Each column represents a genetically distinct mouse strain, and each row corresponds to a specific behavioral measure including locomotion (turning left/right), exploration (rearing), self-directed behaviors (grooming, scratching), and escape responses. Color intensity indicates deviation from the population mean, with red representing increased behavioral expression and blue representing decreased expression relative to the strain average. Z-score thresholding (|Z-score| *>* 1) was applied to all behavioral measures, with escape behaviors displayed separately using modified thresholding parameters to preserve detection of outlier strains exhibiting rare but phenotypically important escape responses. Behavioral measures are stratified by time point (T5, T20, T55) to capture temporal dynamics of phenotypic variation across genetic backgrounds.

In addition to phenotypic diversity due to genotype, we explored sexual dimorphism in our dataset with these new classifiers. We examined the impact of sex on the aggregated phenotype in various strains using a univariate approach. To test for the statistical significance of the effect of sex, we utilized a nonparametric rank test and correcting for multiple testing using false discovery rate (fdr) using Benajamini-Hochberg method. The LOD scores and effect sizes are presented in Figure S7B, with the left panel showing the strength of evidence against the null hypothesis of the non-sex effect. The right panel presents a representation of the direction and magnitude of the effect size with the color and size of circle represents the direction and magnitude of the effect, respectively. The strains highlighted in pink exhibit a significant sex effect for at least one of the aggregated phenotypes. It is important to note that we are generally underpowered with five animals of each sex. However, we find that a high proportion of phenotypes show a sex effect.

### 2.5 JABS-AI (Analysis and Integration) module: Heritability, genetic correlation, and GWAS analysis

Next, one of our goals is to understand the genetic architecture that governs the complex behavioral phenotypes. The quantitative-genetic analysis described in this section provides the backbone for the JABS-AI web platform that follows in section 2.6. The web interface lets any user compute behavioral predictions and downstream genetic analysis for newly uploaded classifiers.

To facilitate this, we utilized the data derived from 49 inbred strains along with 11 F1 hybrid strains, to perform a genome-wide association study (GWAS). It was deemed necessary to exclude the six wild derived strains due to their pronounced divergence, which carried the risk of distorting the outcomes of our mouse GWAS. We first carry out power analysis for both the strain survey datasets (JABS600, JABS1200) using simulation algorithm as proposed by Genome-wide Efficient Mixed Model Association (GEMMA) software. GEMMA, a useful tool for this type of analysis, accounts for population structure and genetic relatedness between individuals, making it ideal for our inbred and hybrid strains. The power analysis as shown in Fig. 8A revealed that we had sufficient statistical power to detect genetic associations. Notably, the JASB1200 dataset demonstrated higher power in detecting these associations compared to the JABS600 dataset. With JABS1200 established as our dataset of choice for conducting the GWAS, we moved forward with assessing each of the 72 phenotypes for their potential association with genotype. We employed the GEMMA software for this purpose, giving particular emphasis to the Wald test p-value in our analysis. These 72 phenotypes have been derived from eight basic classifiers, which include turn left and turn right (each assessed by two different annotators), grooming, scratching, supported rearing, and unsupported rearing. Each of these classifiers has been further categorized into three bout-based measures: average bout length, total duration, and total number of bouts. These bout-based measures were then dissected into three time-based measures (5 minutes, 20 minutes, and 55 minutes) to provide a comprehensive analysis. We tested a substantial number of SNPs (211,077) which necessitated accounting for the inherent correlations among SNP genotypes. To establish an empirical threshold for the p-values, we shuffled the values of one normally distributed phenotype (*TL*_*T* 20_*duration*) randomly and identified the smallest p-value from each permutation. This rigorous process allowed us to set a p-value threshold of 1.9e-05 that reflects a corrected p-value of 0.05. We first report the heritability estimates for phenotypes corresponding to 55 minutes of observed behavior as shown in Fig. 8B. Most of the phenotypes have heritability in the range (0.2-0.8) with bout length based phenotypes having lower heritability relative to bout number or bout duration based phenotypes. Next, to further shed light on the pleiotropic action of genes, we estimate the genetic correlations across these phenotypes using the bivariate linear mixed model implemented in GEMMA. We plot the genomic restricted maximum likelihood (GREML) estimates of bivariate genetic correlations in Fig. 8C. The magnitude of the genetic correlation estimate provides an estimate of genetic overlap (common genetic loci) between two traits, whereas the sign determines the direction of the effects of the overlap on the two traits, i.e., a negative sign corresponds to the effect in the opposite direction on the two traits and vice versa. We hypothesize that for a given behavior, the bout-based measures within the behavior share common genetic effects and affect the traits in the same direction. Indeed, we find estimates of genetic correlations that are positive between the number of bouts (nBouts) and duration of the behavior (duration), the average length of each bout (avgLen) and duration of the behavior (duration), and the number of bouts (nBouts) and duration of the behavior (duration) for all behaviors except the turn right behavior (A1_TR) by annotator 1. We find positive estimates of genetic correlations between two annotators (A1, A2) for the turn left or right behaviors since we expect the genetic architecture underlying the same behavior from two annotators to overlap maximally.

**Figure 8:**
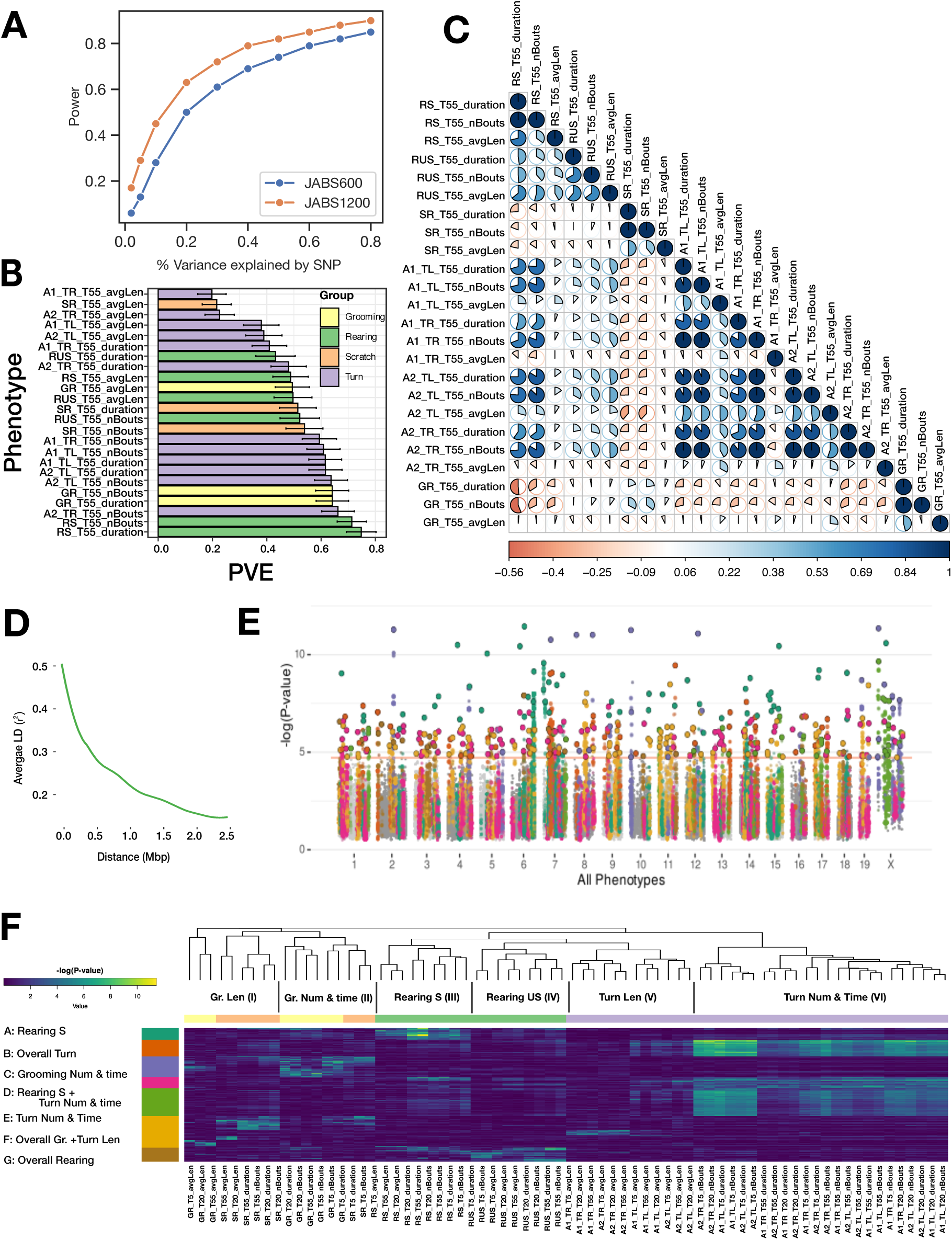
JABS-AI (Analysis and Integration) module: Large-scale GWAS investigation of different mouse behaviors utilizing the JABS1200 dataset. : (A) Statistical power comparison between two datasets (JABS600 vs JABS1200) at the genome-wide significance threshold of 2.4e-07. The y axis shows how power varies with SNP effect size (x axis) (B) Aggregate (55 min) phenotypes’ heritability (PVE) estimates. (C) Lower Triangular Matrix Representation of Genotypic Correlation Among all of the 55 minute aggregate phenotypes using a bi-variate linear mixed model, (D) Linkage disequilibrium (LD) blocks size, along with the mean genotype correlations for SNPs at varying genomic distances. (E) Aggregated GWAS results graphically represented via a comprehensive Manhattan plot. Peak SNP clusters, extracted from (F), determine color differentiation; SNPs within the same LD block are color-coordinated to match their peak SNP. Each SNP is assigned the minimum p-value derived from all phenotypes. (F) An inclusive heatmap exhibiting all the significant peak SNPs for each phenotype. Each row, representing an SNP, is color-coordinated according to the allocated cluster within the k-means clustering. The color scheme originating from the k-means cluster is also applied in panel E of this analysis.

We adopted a specific approach to identify quantitative trait loci (QTL): we started with the SNP that exhibited the lowest p-value across the genome and designated it as a locus. We then grouped together adjacent SNPs showing a significant level of correlation in their genotypes (*r*^2^ ≥ 0.2), employing a greedy strategy. We continued this process, moving on to the next SNP with the lowest p-value until we allocated all significant SNPs to a QTL. Given the inherent genetic structure of inbred mouse strains, large linkage disequilibrium (LD) blocks are expected, as represented in Fig. 8D.

Additionally, we observe pleiotropy with certain loci displaying significant associations with multiple phenotypes, an anticipated occurrence given the correlation among many of our phenotypes and the potential for individual traits to be influenced by similar genetic loci. To get a clearer picture of the pleiotropic structure apparent in our GWAS findings, we constructed a heatmap (fig. 8F) of significant QTL across all phenotypes and employed *kmeans* clustering to identify QTL sets governing phenotype groups. The phenotypes are grouped into 6 categories namely: grooming bout length, grooming bout number and total time, rearing supported, rearing unsupported, turn bout length, turn bout number and total time. We uncovered seven unique clusters of QTLs (A-G), each regulating a different combination of these phenotype subgroups (fig. 8F). Clusters B and G notably held pleiotropic QTLs that influenced overall turn and rearing behaviors, respectively. Yet within cluster F, we identified distinct QTL sets - one that steered grooming behavior, and another, non-overlapping set that determined turn length. This distinction signifies the existence of distinct genetic underpinnings for these different behaviors even within the same cluster. Finally, we color the associated SNPs in the Manhattan plot (fig. 8E) showing QTLs associated with all phenotypes.

### 2.6 Data Integration: A web application for classifier sharing and downstream genetic analysis

In conjunction with the release of the curated datasets and the JABS active learning GUI app (JABS-AL), we have developed and launched a web-based application, JABS-AI (analysis and integration), aimed at streamlining the sharing and utilization of classifiers. Through this platform, users can view, download and rate the classifiers for various behaviors that have been developed and trained in our laboratory as shown in Figure 9B. In addition, it provides an insight into their heritability scores and offers a feature to examine the pair-wise genetic correlations amongst different phenotypes. An added functionality of this web application is that it allows users to upload their own classifiers (trained using JABS-AL GUI app) for any specific behavior. Upon uploading, the application automatically executes the classifier on a dataset of the user’s choosing from our strain survey datasets. It conducts an automated analysis of behavior and genetics, and subsequently dispatches the results to the user’s designated email address within a few hours (see figure 9A). The results coming out of webapp would contain following downstream analysis on the dataset selected by the user in the app:

1. Density plots of predicted behavior (similar to Figure 5D,G).
2. Strain-behavioral phenotype heatmap (similar to Figure 7).
3. Heritability and genetic correlation of behavioral phenotypes (similar to Figure 8B,C).

**Figure 9:**
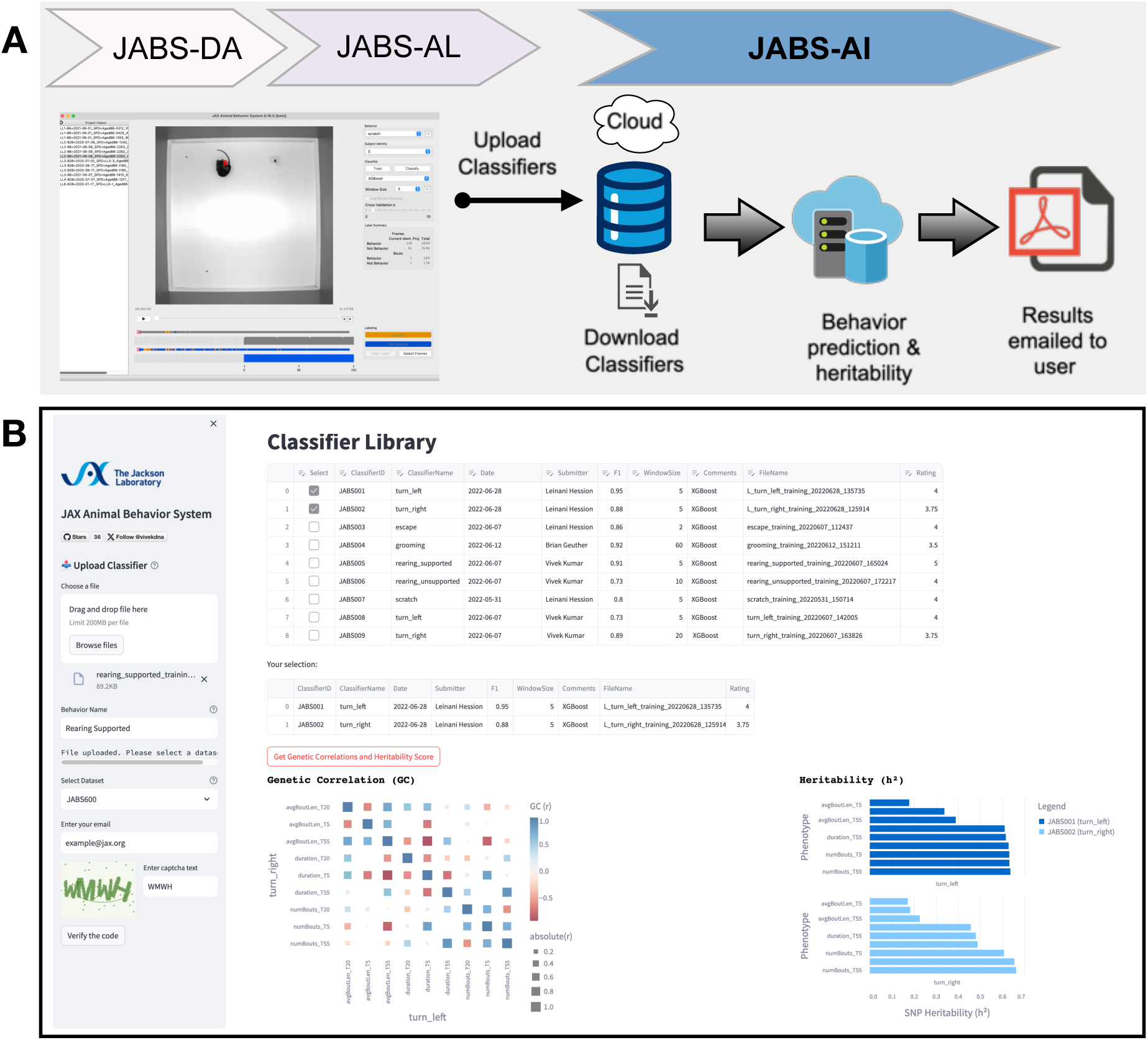
**JABS-AI (Analysis and Integration) : A web application for sharing the JABS classifiers and automated downstream genetic analysis**: (A) Illustrates the fundamental workflow of the web application, beginning with the user employing a classifier trained via the JABS active learning application. The user subsequently deposits this classifier into our web application, which performs comprehensive automated analyses, encompassing both behavioral and genetic aspects, on dataset selected by the user from our curated strain survey collection (JABS600, JABS1200) accessible via a dropdown menu. The outcome of these analyses, encapsulating detailed behavioral patterns and genetic correlations, are then dispatched to the user’s designated email address within a short timeframe. (B) Screenshot of the webapp highlighting the tabular presentation of the repository of classifiers developed in our laboratory, complete with pertinent metadata such as the date of creation, training hyperparameters, and user ratings. When any two classifiers are selected, the application offers the option to analyze the genetic correlations between the phenotypes corresponding to the selected classifiers, in conjunction with their heritability scores.

This web application serves as a facilitative tool aimed at fostering collaboration among researchers and streamlining the advancement of automated behavior quantification studies by providing a platform for the efficient sharing and analysis of behavioral classifiers.

## 3 Discussion

Democratization of machine vision methods for advanced behavior quantification remains a challenge. Often, tracking and behavior classifiers are not transferable between laboratories. This limits the reuse of prior work with each laboratory essentially starting from scratch with advanced behavior quantification. JABS and the companion DIV Sys are designed to overcome these limitations. JABS components include video data acquisition, behavior annotation, classifier sharing, and genetic analysis. By adopting the JABS-DA laboratories can use our pose estimation and segmentation models that work across 62 mouse strains of varying coat colors and sizes. This greatly eases the barrier to entry for advanced behavior quantification. The next steps of creating behavior classifiers are carried out using JABS-AL, an active learning system modeled after JAABA [21]. We benchmarked JABS-AL using grooming data set and demonstrate that it reaches very good performance with 10% of the data needed for a 3D-CNN for action detection. Once constructed, behavior classifiers can be shared through JABS-AI, a cloud based tool. Labs can create their own behavior classifiers using JABS-AL or download an existing one for use from another lab from JABS-AI in order to annotate single behaviors. The power of JABS-AI is also in the embedded strain survey data. A deposited classifier is inferred on one of three datasets and heritability and genetic correlation results are returned.

A key decision point is the adoption of common apparatus to create a uniform visual look of the video data across laboratories. This enables cross application of foundational models and exchange of behavior classifiers across labs. We realize that this may be challenging for some labs with limited space and budget. Indeed, JABS has a large footprint with a 2×2×6 feet (W x L x H) space requirement, costs several thousand dollars for components, and requires some computational expertise to set up and operate. Laboratories must balance this cost with the labor and time costs of adopting an existing set-up for advanced behavior analysis. In lieu of adopting a common apparatus, efforts are being made to build foundational models that can handle diverse environments and even animals [49]. While similar datasets are common in human pose estimation, the development of equivalent datasets for animals is still underway. However, such foundational models are not yet available. Even when they are available, the initial abstraction step is simple compared to the later step of behavior classification. For instance, in our gait and posture paper, producing a pose estimation model that generalizes to diverse mouse strains took approximately 6 months. It was an iterative hard example mining task. However, the process of deriving gait and posture from keypoints and its genetic validation took over 1.5 years. By simply adopting JABS, laboratories gain access to both the pose model and validated gait/posture algorithms.

While our pose estimation model was not specifically trained on tethered animals, published research demonstrates that keypoint detection models maintain robust performance despite the presence of headstages and recording equipment. Once accurate pose coordinates are extracted, the downstream behavior classification pipeline operates independently of the pose estimation method and would remain fully functional. We recommend users validate pose estimation accuracy in their specific experimental setup, as the behavior classification component itself is agnostic to the source of pose coordinates.

Another added benefit to JABS is the ability to apply novel behavior classifiers to large-scale genetically diverse datasets collected at JAX through JABS-AI. We have modeled this after existing platforms such as GeneNetwork and DO QTL. Currently, we provide heritability estimates of any classifier that is deposited. Users can also select behaviors to genetic correlation studies. Thus, even if two behaviors are different, they may measure the same genetic architecture. Although the current version of JABS-AI does not offer GWAS analysis due to compute restrictions, the method can be easily extended for such analysis. It is also feasible to link animal behaviors to human traits through PheWAS analysis [8, 50]. This would provide even more detailed information for users about the genetic regulators of complex behaviors. The current data sets, JABS600, JABS1200, and JABS-BxD consist young wild type animals. We have collected datasets in aging populations with various frailty statuses and animals that display nocifensive behaviors. These could also be integrated into JABS-AI for preclinical behaviors independent of genetic analysis.

While JABS is designed for individual behavior annotation, a common task in behavioral neurogenetics to determine an internal state, e.g. anxiety or social state. Often these are accomplished by measuring a single behavior. A more powerful approach is the application of behavior indices to predict certain states. These indices can be constructed using multiple behaviors and even other covariates. For example, we trained a model to predict frailty using data from over 600 JABS-DA open field tests from C57BL/6J mice of varying age and frailty. We used 34 features from JABS to derive frailty. Similarly, in companion work, we derived pain scale with 82 features from JABS. These indices were constructed with almost 1000 animals and can readily be transferred to other labs that collect data using the JABS system. This is incredibly powerful and allows labs to leverage each other’s models by using a common platform. Similarly, for pain states, we have tested multiple strains and built a pain intensity models that can be utilized. We believe a true advantage of advanced phenotyping using video data is the ability to reuse and extract more information from existing data. This essentially allows us to use less animals, a core 3R principle.

Here, we describe JABS as a single animal open-field assay that lasts from minutes to a few hours. However, we have designed the JABS arena for long-term housing of animals with a food hopper and lixit. This was the primary reason we worked with JAX-IACUC to certify JABS-DA for long-term monitoring with key required environment measures. By blocking visible light and imaging using IR LED illumination, we obtain uniform data at night or day. We routinely collect video data with three mice over several days. The models for tracking, instancing, and identity maintenance need to evolve. We plan to extend JAB-AI with classifiers for social interactions and homeostatic behaviors. Thus, future iterations of JABS will develop and share multi-animal behavior analysis.

Even when data acquisition is standardized, another fundamental source of variability can enter the system when different human experts within or across different labs, annotate the same videos for the same behavior. This type of variability can arise due to variety of factors, including differences in training, personal biases, and individual interpretation of behavior. As behaviors become more complex, we expect behaviorists to show more disagreement. These disagreements could be as simple as varying understanding of the starts and stops of the behaviors. Or more fundamentally differing opinion on the behavior such as between aggression and play. In a previous study, we asked 5 humans to annotate grooming behavior and found that the agreement ranged from 86% to 91%. In this case we simply compared on a frame wise basis labels from annotators who were asked to label every frame in the same set of videos. In most cases, such comparisons are infeasible. A more realistic comparison is provided in this manuscript. Two annotators built their own classifiers for left and right turn behaviors. Then we compared the predictions from each set of classifiers on JABS-DA video. This is akin to two different laboratories that may deposit classifiers for the same behavior. JABS-AI does not save primary the training data from each annotator (lab), and simply uses the trained classifiers to infer on a new set of videos (e.g. JABS600). From these we can compare the overlap between the two classifiers’ inferences. We do not make assumptions about which classifier is the ground truth and simply compare both classifiers.

In section 2.4.1, we demonstrate that even for simple behaviors like left and right turn, there is a significant amount of disagreement between predictions coming from classifiers trained by two expert annotators within the lab for the same behavior. One of the most commonly used statistical measures to quantify the inter-annotator variability is Cohen’s kappa which assesses the level of agreement between the annotators taking into account the possibility of agreement by chance. The Cohen’s kappa statistic works well for frame-wise comparison but is ill-defined for bout-wise comparison as unlike frames, bouts are not conserved. In order to overcome this limitation, we have introduced a new approach based on graph theory, called the *ethograph*. This network approach allows us to define measures that quantify the agreement between two annotators when comparing bouts of behavior among different annotators. By comparing the entire sequences of frames, the ethograph reduces subjectivity and allows for a more holistic and consistent interpretation of behaviors. This makes it well-suited for bout-wise comparison, and may provide a more accurate estimate of inter-annotator agreement than the frame-based kappa statistic.

Even though frame wise comparison shows the overlap performance is poor (κ = 0.64 and 0.65 for Left and Right Turn, respectively), each classifier does a good job of identifying tuning behaviors. The turn behaviors are short in length of frames, and the two annotators differ in defining the starts and stops of the behaviors. One annotator labels just the core turn behavior, and the other starts labeling turning behavior a few frames earlier and ends later. Classifiers from both annotators generally find the same bouts of turning which we could visualize in the ethograph. We explored bout-wise accuracy metrics as an alternative to frame-wise metrics. We also explore post-processing predictions using hyperparameters for filter and stitch. By adding these, we observe much higher agreement between the classifiers from two annotators for the same behaviors (overlap increases from 49% to 61%). It is important that users clearly define the behavior as best as possible, and document the filter and stitch parameters.

JABS users, when confronted with multiple classifiers for the same behavior in JABS-AI, must prioritize use of one classifier. JABS-AI offers a genetic solution to this challenge - by prioritizing the classfier that is more heritable. Heritability is an estimate of variance explained by genetics and can act as a discriminator in this situation. We also calculate genetic correlation with allows users to determine which underlying genetic construct is being measured. For instance, both left and right turn are highly genetically correlated and, therefore for the purposes of genetics, there is simply turn behavior. However, for certain unilateral models such as brain lesions, stroke, optogenetic stimulation, or injury, the ability to distinguish left and right turns can be critical.

### 3.1 Supervised vs unsupervised behavior segmentation

Unsupervised methods like Keypoint-MoSeq [51], VAME [52], B-SOiD [22], and MotionMapper [53] which prioritize motif discovery from unlabeled data but may yield less precise alignments with expert annotations, as evidenced by lower F1 scores in comparative evaluations [54]. Supervised approaches (like ours), by contrast, employs fully supervised classifiers to deliver frame-accurate, behavior-specific scores that align directly with experimental hypotheses. Ultimately, a pragmatic hybrid strategy, starting with unsupervised pilots to identify motifs and transitioning to supervised fine-tuning with minimal labels, can minimize annotation burdens and enhance both discovery and precision in ethological studies.

### 3.2 Future directions and challenges

We see several areas of improvement in JABS in the future. First, the success of such a platform depends on community adoption. As such, JAX has made JABS free for noncommercial use, and we have listed all parts and software used to make JABS. We realize that many laboratories may not have the computational or fabrication resources to construct JABS and that commercial suppliers who can provide a turnkey system are needed for JABS-DA. JABS-AL and JABS-AI require fewer, though still significant, resources to support.

JABS-AI currently does not support upload of training videos due to resource limitations. This prevents other users from interrogating the primary training labels. It also prevents users from downloading and labeling new behaviors or modifying classifiers that have been uploaded. Future versions could support sharing of complete training data instead of the classifier only.

Furthermore, since the classifiers are trained on few densely labeled short video recordings and then further make predictions on a large strain survey consisting of multiple strains of mice, there is some variability in predictions purely due to out-of-distribution strains in the strain survey. Therefore, the inter-annotator variability in predictions on the new set of strains of mice can be attributed to both the variability in the human labeling and genetic variability in the strain survey. Calculating the heritability scores might help in this scenario by providing us a quantitative measure of the extent to which the inter-annotator variability is due to genetic factors versus interpretation by the human labelers.

#### 3.2.1 Rodent Homes and Hotels

Finally, JABS and DIV Sys are complementary systems that enable behavioral monitoring across multiple scales and resolutions. DIV Sys facilitates long-term observation of home-cage behaviors, whereas JABS offers high-resolution tracking of gait, posture, and other discrete actions [9, 10]. The larger space in JABS can potentially accommodate additional tasks designed to probe specific neural circuits [55, 56], and neural recordings can be collected from instrumented mice in this same arena. We see these as "ethological tasks" that can be performed continuously over long-periods of time in order to interrogate neural and genetic circuits in customizable environments – "hotels". Examples include mazes and other tasks that neurobehavior researchers have been developing. These assays can be validated using genetic or pharmacological models on a shared platform such as JABS. These two platforms provide a dual approach: continuous surveillance of mice in their home-cage environments (via DIV Sys) alongside targeted assessments of particular behaviors in a dedicated hotel arena (via JABS). This combined paradigm presents a powerful framework to link genetic and neural changes to complex behaviors. Indeed, elucidating how altered behaviors result from altered neural circuits and altered genetic pathways remains a central challenge in computational ethology one that platforms such as JABS and DIV Sys are poised to address.

## 4 Materials and Methods

### 4.1 Animals

All animals were obtained from The Jackson Laboratory production colonies or bred in a room adjacent to the testing room as previously described [8, 9, 24]. All behavioral tests were performed in accordance with approved protocols from The Jackson Laboratory Institutional Animal Care and Use Committee guidelines.

### 4.2 Datasets

To facilitate reproducibility and community-driven discovery, we have made three comprehensive datasets publicly available. Each dataset contains multiple open-field arena (OFA) videos (one hour recording per video) and corresponding pose-estimation keypoint files.

#### JABS600

This dataset comprises approximately 600 videos from 62 genetically diverse mouse strains, with sexes balanced within each strain. It serves as a broad survey for initial explorations of behavioral phenotypes across a wide genetic landscape. https://doi.org/10. 7910/DVN/SAPNJG

#### JABS1200

An extension of the first dataset, JABS1200 contains nearly 1,200 videos from the same 62 strains, effectively doubling the sample size per strain. This increased depth provides greater statistical power for detecting significant associations in Genome-Wide Association Studies (GWAS). https://doi.org/10.7910/DVN/SAPNJG

#### JABS-BxD

This collection includes over 1,000 videos from 108 BxD recombinant inbred mouse strains, which are derived from a cross between the C57BL/6J (B6) and DBA/2J (D2) parental strains. With approximately five males and five females per strain, this dataset is structured to support high-resolution genetic mapping of behavioral traits. https://doi.org/10.7910/DVN/RQYI04

#### JABS behavioral classifier example projects

Complete training resources for behavioral classification, comprising experimental video data, extracted pose coordinates, and corresponding behavioral annotations, are made publicly available. https://doi.org/10.5281/zenodo.16697332

### 4.3 JABS workflow guide

The JAX Animal Behavior System (JABS) provides an integrated, four-stage pipeline guiding users from standardized data acquisition to novel genetic discovery. This workflow utilizes three core software modules (JABS-DA, JABS-AL, JABS-AI) and a standalone pose estimation engine, as detailed in the summary below.

| JABS End-to-End Behavioral Phenotyping Workflow |  |  |
| --- | --- | --- |
| Step | Module & Objective | Resources |
| **1. Data Acquisition** | JABS-DA: Capture uniform, high-fidelity video recordings to ensure cross-laboratory reproducibility. | Data acquisition pipeline |
| **2. Pose Estimation** | Mouse Tracking Runtime: Convert video into a quantitative representation of posture by tracking 12 body keypoints. | Tracking Runtime pipeline |
| **3. Behavioral Annotation & Classifier Training** | JABS-AL: Train supervised classifiers for user-defined behaviors using a sparse-labeling approach. | JABS-AL GUI app<br>JABS-AL Tutorial |
| **4. Genetic Analysis** | JABS-AI: Deploy classifiers on large-scale genetic datasets to discover genetic drivers of behavior. | JABS-AI Webapp |

### 4.4 Pose tracking pipeline

We have previously published our single mouse pose model here [10], with the training data and trained models available at https://zenodo.org/records/6380163. We have also released our pose tracking pipeline, which includes single and multimouse tracking and classifier prediction at https://github.com/KumarLabJax/mouse-tracking-runtime. In addition to our inhouse tracking pipeline, we have made our pose format accessible via SLEAP-io conversion hooks (https://github.com/talmolab/sleap-io).

### 4.5 Grooming benchmarking study

The Convolutional neural network (CNN) applied to the grooming benchmark dataset follows a typical feature encoder structure except using 3D convolution and pooling layers instead of 2D. The final layer was used as the output probabilities for not grooming and grooming predictions for each frame. The exact architecture and the training details are described in detail here [9]. Furthermore, the JABS grooming classifier has been trained with both XGBoost [37] and Random Forest model [38] with their default hyper-parameters in the XGBoost and scikit-learn library respectively.

To assess classifier performance on frame-by-frame labeling of animal behavior, we report five standard metrics:

#### Accuracy

Percentage of all video frames correctly labeled as either target behavior or not.

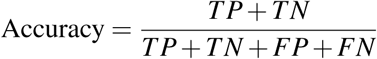

#### F1 score

Harmonic mean of precision (fraction of predicted target frames that are correct) and recall (fraction of actual target frames detected).

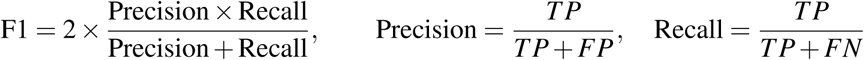

#### AUROC

Area under the receiver-operating-characteristic curve; measures how well the classifier separates frames containing the target behavior from all others, independent of the decision threshold (k).

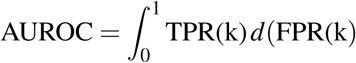

#### True Positive Rate (TPR)

Proportion of frames with the target behavior that the classifier correctly labels.

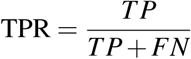

#### False Positive Rate (FPR)

Proportion of frames without the target behavior that are incorrectly labeled as target by the classifier.

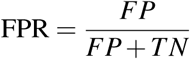

#### TPR@5 % FPR

In practical applications, it is important to capture as many true instances of the behavior as possible while avoiding excessive false positives, which reduce trust in the predictions and increase validation effort. Therefore, we also report the true positive rate (TPR) achieved when the classifier is tuned to allow at most 5% false positive rate (FPR).

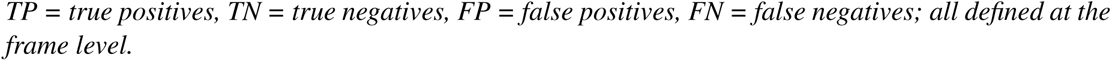

### 4.6 Downstream analysis on strain survey

#### 4.6.1 Aggregate phenotypes

For each classifier, we construct nine aggregate phenotypes corresponding to the three metrics (total duration, number of bouts and average bout length) and three time bins (first 5, 20 and 55 minutes). For instance, a phenotype named “turn_left_T55" (in Figure 7) represents the total duration of left turn behavior in the first 55 minutes of the video averaged across all videos. For more details, refer to supplementary tables S2 and S3.

#### 4.6.2 Z-score normalization of behavioral phenotypes

To facilitate comparison across strains and behaviors, each phenotype was standardized using z-score normalization. This process transforms the raw phenotype values into units of standard deviation relative to the mean across all strains. Specifically, for each phenotype *x*, the z-score *z* is calculated as

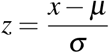

where µ is the mean and σ is the standard deviation of the phenotype values across all strains. This normalization centers the data around zero and scales it so that the spread reflects variability within the dataset. In our heatmap (Figure 7), colors correspond to z-scores, emphasizing how much a strains behavior deviates above (red) or below (blue) average. To improve clarity, we highlight entries with |*z*| *>* 1, indicating behaviors differing from the mean by more than one standard deviation.

#### 4.6.3 Genetic analysis

The aggregate behavioral phenotypes were analyzed to study the genetic associations of strain-specific behaviors. Genotype data for the different mouse strains were obtained from the Mouse Phenome Database (https://phenome.jax.org/genotypes). We used genotypes derived from the Mouse Diversity Array (MDA), where di-allelic genotypes were inferred from parental genomes. Quality control filters retained SNPs with a minor allele frequency (MAF) of at least 10% and a maximum of 5% missing data.

Genome-wide association studies (GWAS) were conducted using the R package mousegwas, as previously described in [9]. Classical laboratory mouse strains were included in the analysis, excluding wild-derived strains. Associations were computed using the linear mixed model (LMM) method implemented in GEMMA. To reduce confounding from linkage disequilibrium near tested markers, a Leave One Chromosome Out (LOCO) approach was applied: for each chromosome under test, the kinship matrix was calculated using SNPs from all other chromosomes.

To control for multiple testing and establish an appropriate genome-wide significance threshold, we performed permutation-based empirical calibration. Specifically, phenotype values from a randomly sampled continuous trait were shuffled across individuals multiple times, and the minimum p-value from each permutation was recorded. This approach yielded a p-value threshold of approximately 1.9*e* − 05, corresponding to a family-wise error rate of 0.05 after correcting for the number of tests performed.

SNP-based heritability for each behavioral phenotype was also estimated using mousegwas. For each phenotype, a genetic relatedness matrix (GRM) was constructed from quality-controlled SNPs (filtered for MAF and missingness). The analysis was performed on the JABS1200 dataset, comprising 1,139 individuals and 211,070 SNPs. Fixed-effect covariates included sex, weight, and coat color of the animals.

The GWAS execution was wrapped in an R package called mouseGWAS available on github: https://github.com/TheJacksonLaboratory/mousegwas.

For genetic correlation analyses, we applied GEMMA’s bivariate LMM (using this pipeline: https://github.com/gautam-sabnis/genetic_correlation) to each pair of phenotypes.

#### Power analysis simulation for GWAS

To assess the statistical power of our GWAS design, we followed the simulation-based approach described in the original GEMMA paper [57]. Briefly, we first filtered SNPs from the GEMMA output (for a randomly selected behavioral phenotype) to retain only those with nominal association (*p <* 0.05), sorted them by genomic position, and then selected a fixed number of evenly spaced SNPs as "causal." For each causal SNP, we assigned an effect size r<u>equired to explai</u>n a specified proportion of phenotypic variance (PVE), calculated as 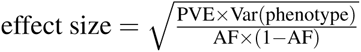, where AF is the SNP allele frequency. We simulated new phenotypes by adding the effect of each causal SNP to the original phenotype, re-ran GWAS using GEMMA, and defined power as the proportion of simulated causal SNPs detected above the genome-wide significance threshold (e.g., Bonferroni-corrected *p <* 0.05).

## 5 Acknowledgments

We thank members of the Kumar Lab for helpful advice and Leinani Hession for training behavior classifiers. Michelle Foskett (Process Quality Control) and Rosalinda Doty (Diagnostic and Pathology Services) help with environment and pathology data. This work was funded by The Jackson Laboratory Directors Innovation Fund, National Institute of Health DA041668 (NIDA), DA048634 (NIDA), MH138309 (NIMH), and AG078530 (NIA). All code and training data will be available at Kumarlab.org and Kumar Lab Github (https://github.com/KumarLabJax).

## 6 Supplementary Material

### 6.1 Grooming benchmark dataset

**Table S1:** Data used for grooming benchmark. Number of videos (first column), and number of annotated frames (second and third column).

| Video Count | Annotated Frames |  |
| --- | --- | --- |
|  | Behavior | Not Behavior |
| 10 | 11778 | 18201 |
| 20 | 19540 | 28777 |
| 50 | 43022 | 51466 |
| 100 | 76109 | 104889 |
| 500 | 399155 | 595798 |
| 1100 (All) | 863394 | 1318396 |

**Table S2:** Summary of framewise behavioral phenotypes and their definitions. Each value corresponds to the total duration (in seconds) of the indicated behavior during the specified time window, averaged across all analyzed videos.

| Phenotype | Description |
| --- | --- |
| turn_left_T5 | Total duration of left turn behavior during the first 5 minutes of video. |
| turn_left_T20 | Total duration of left turn behavior during the first 20 minutes of video. |
| turn_left_T55 | Total duration of left turn behavior during the first 55 minutes of video. |
| turn_right_T5 | Total duration of right turn behavior during the first 5 minutes of video. |
| turn_right_T20 | Total duration of right turn behavior during the first 20 minutes of video. |
| turn_right_T55 | Total duration of right turn behavior during the first 55 minutes of video. |
| grooming_T5 | Total duration of grooming behavior during the first 5 minutes. |
| grooming_T20 | Total duration of grooming behavior during the first 20 minutes. |
| grooming_T55 | Total duration of grooming behavior during the first 55 minutes. |
| rearing_supported_T5 | Total duration of supported rearing during the first 5 minutes. |
| rearing_supported_T20 | Total duration of supported rearing during the first 20 minutes. |
| rearing_supported_T55 | Total duration of supported rearing during the first 55 minutes. |
| rearing_unsupported_T5 | Total duration of unsupported rearing during the first 5 minutes. |
| rearing_unsupported_T20 | Total duration of unsupported rearing during the first 20 minutes. |
| rearing_unsupported_T55 | Total duration of unsupported rearing during the first 55 minutes. |

### 6.2 Quantifying strain survey dataset imbalance

Strain Imbalance (SI):

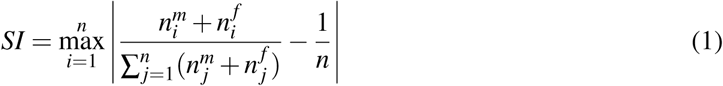

Gender Imbalance (GI) for each strain *i*:

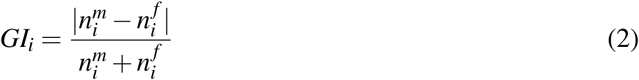

**Figure S1:**
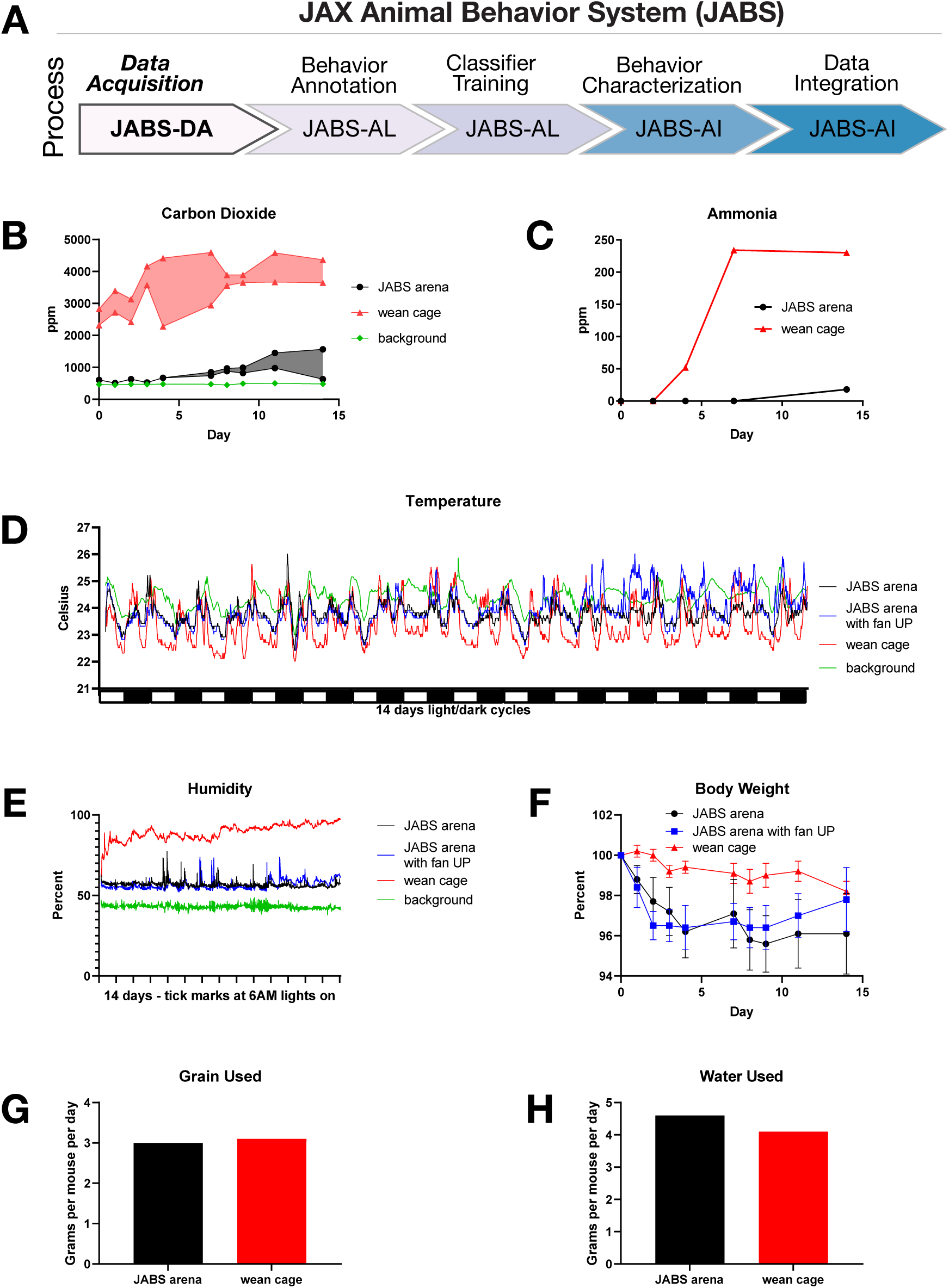
JABS data acquisition module: Environmental parameters in the arena. (A) JABS pipeline highlighting individual steps towards automated behavioral quantification. (B) Carbon dioxide concentrations and (C) ammonia concentrations were both much higher in the standard wean cage than in the JABS arena. Carbon dioxide was also compared to room background levels. (D) Temperature and (E) humidity measured at floor level in JABS arenas and a standard wean cage compared to room background across a 14 day period. (F) Average body weight as percent of start weight in each JABS arena and wean cage across the 14 day period. (G) Food and (H) water consumption shown as grams per mouse per day for one JABS arena and one wean cage for a 14 day period.

**Figure S2:**
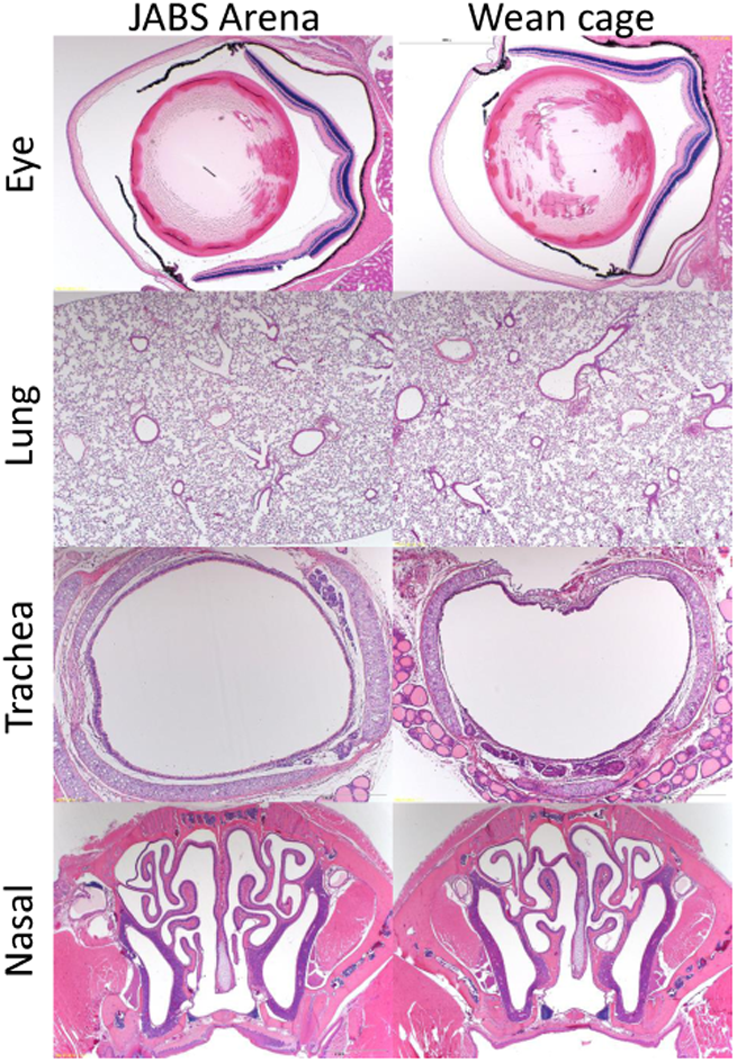
Representative hematoxylin and eosin (H&E) stained tissue sections from mice after spending 14 days in the JABS arena or control wean cage. Tissues selected for examination (eye, lung, trachea and nasal passages) are those expected to be most affected if the mice lived in a space with inadequate air flow. All tissues appeared normal.

The Average Gender Imbalance (AGI) can be calculated as the mean of the Gender Imbalance (GI) for all strains:

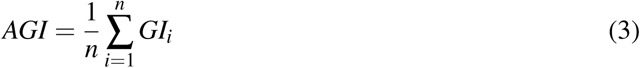

1. *n* is the number of strains
2. 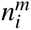 is the number of male samples for strain *i*
3. 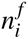 is the number of female samples for strain i.

### 6.3 List of features for JABS

**Table S4:**
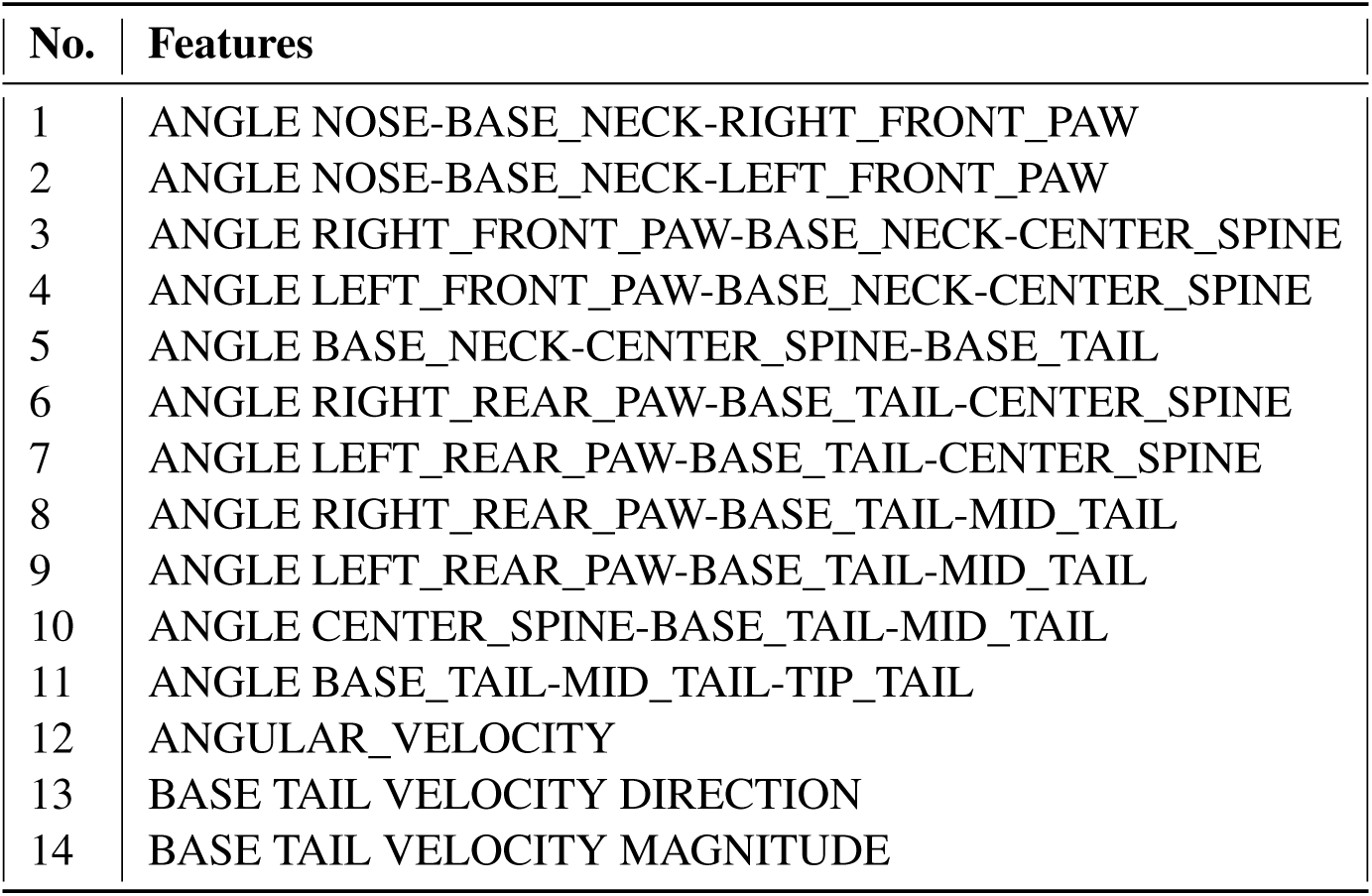
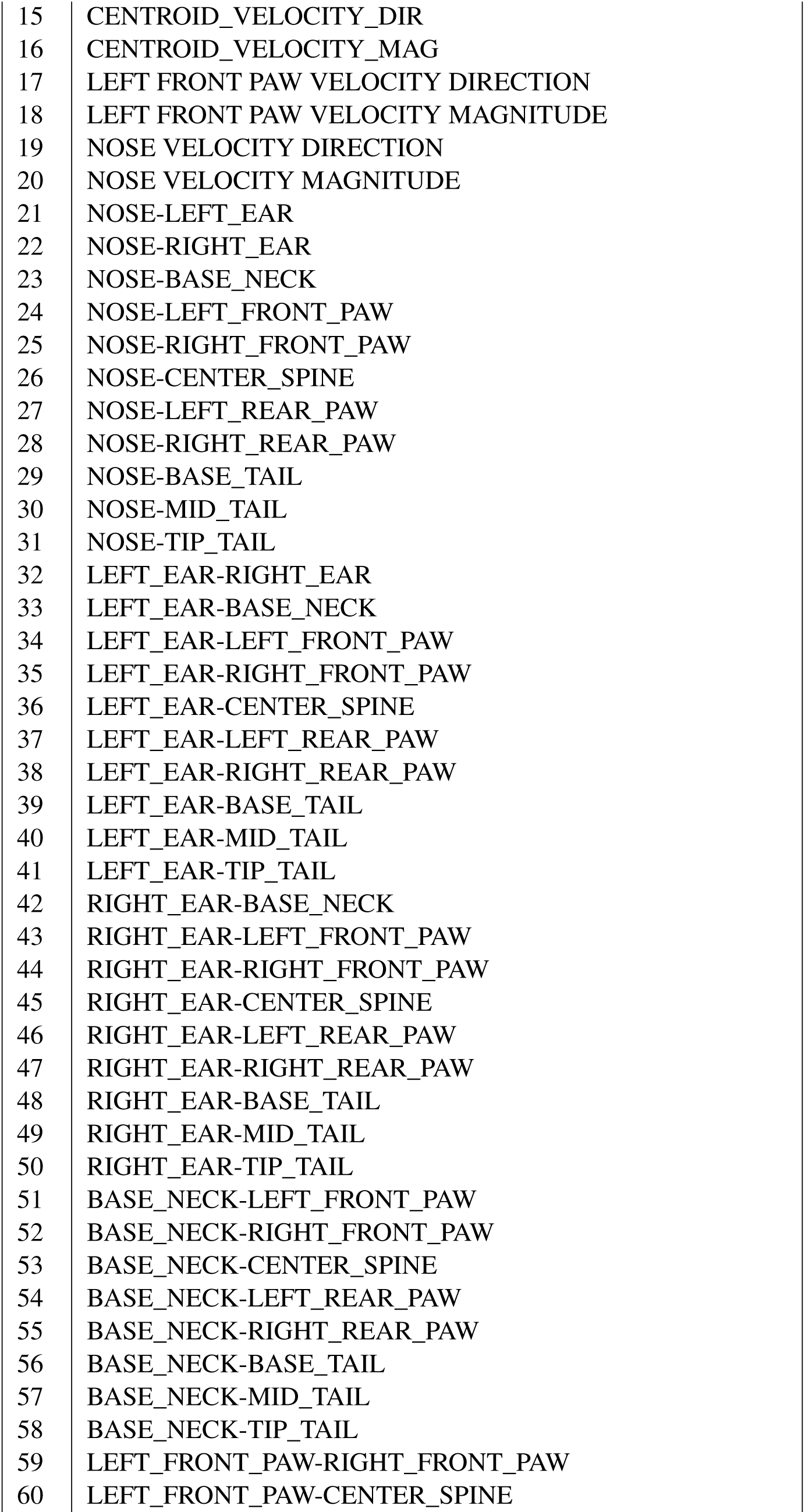
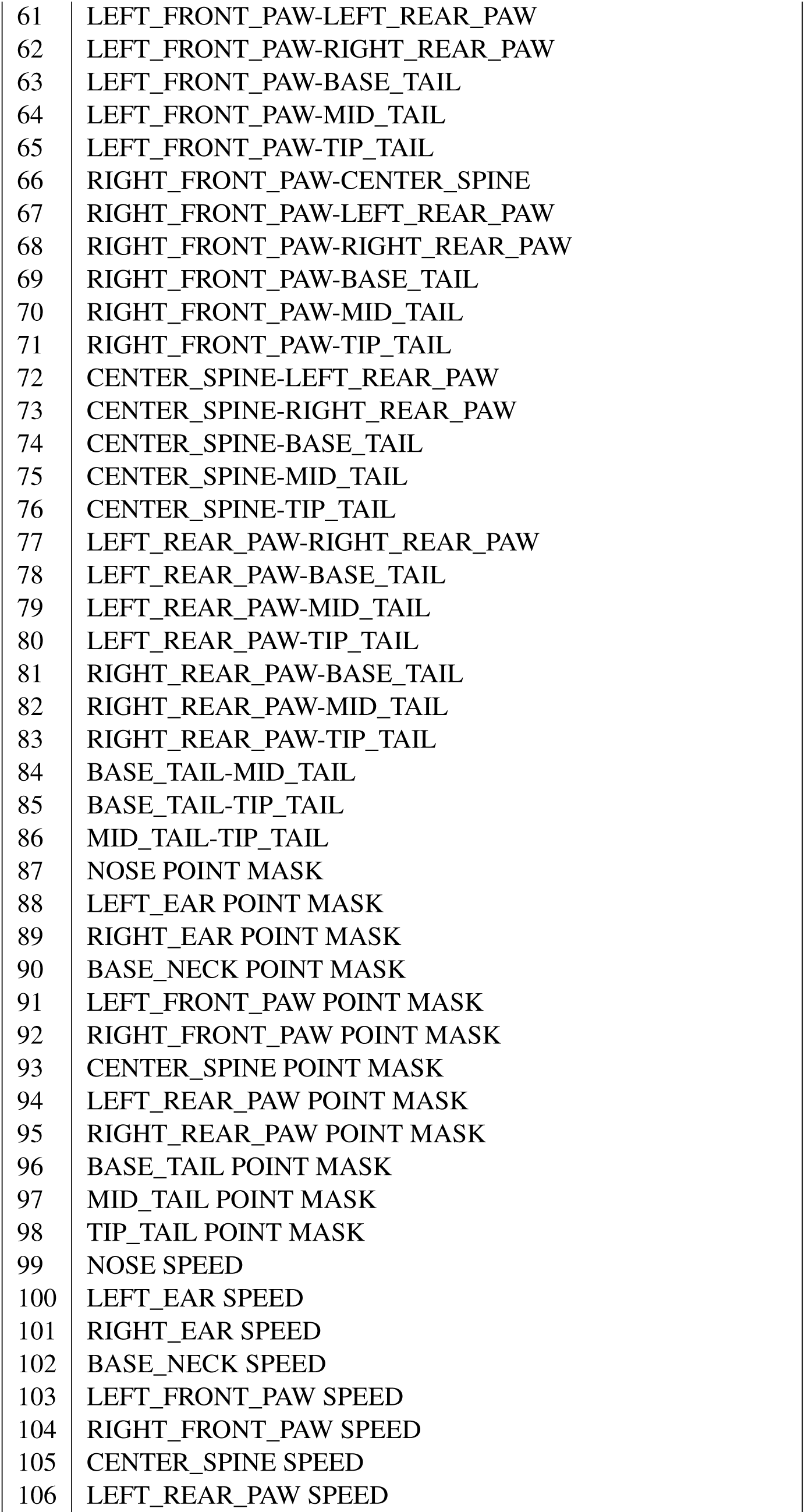
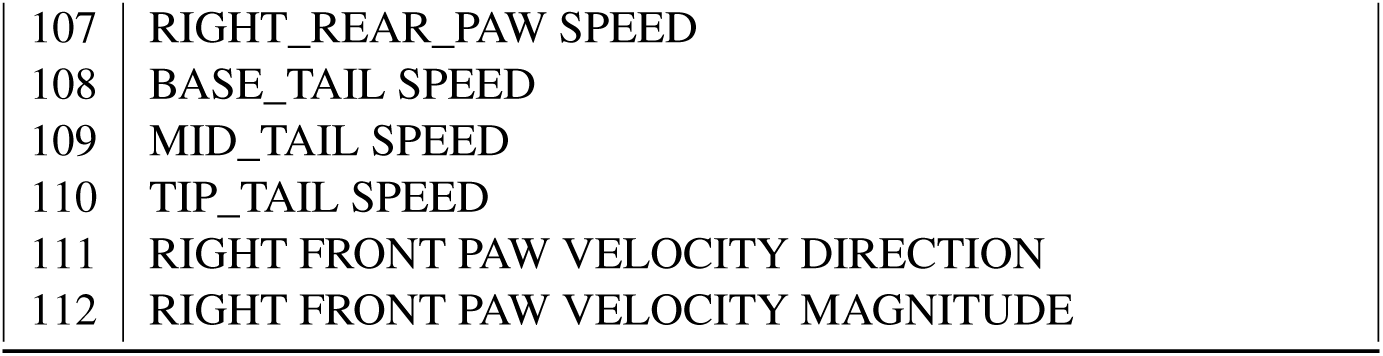
List of JABS features.

| No. | Features |
| --- | --- |
| 1 | ANGLE NOSE-BASE_NECK-RIGHT_FRONT_PAW |
| 2 | ANGLE NOSE-BASE_NECK-LEFT_FRONT_PAW |
| 3 | ANGLE RIGHT_FRONT_PAW-BASE_NECK-CENTER_SPINE |
| 4 | ANGLE LEFT_FRONT_PAW-BASE_NECK-CENTER_SPINE |
| 5 | ANGLE BASE_NECK-CENTER_SPINE-BASE_TAIL |
| 6 | ANGLE RIGHT_REAR_PAW-BASE_TAIL-CENTER_SPINE |
| 7 | ANGLE LEFT_REAR_PAW-BASE_TAIL-CENTER_SPINE |
| 8 | ANGLE RIGHT_REAR_PAW-BASE_TAIL-MID_TAIL |
| 9 | ANGLE LEFT_REAR_PAW-BASE_TAIL-MID_TAIL |
| 10 | ANGLE CENTER_SPINE-BASE_TAIL-MID_TAIL |
| 11 | ANGLE BASE_TAIL-MID_TAIL-TIP_TAIL |
| 12 | ANGULAR_VELOCITY |
| 13 | BASE_TAIL_VELOCITY_DIRECTION |
| 14 | BASE_TAIL_VELOCITY_MAGNITUDE |
| 15 | CENTROID_VELOCITY_DIR |
| 16 | CENTROID_VELOCITY_MAG |
| 17 | LEFT_FRONT_PAW_VELOCITY_DIRECTION |
| 18 | LEFT_FRONT_PAW_VELOCITY_MAGNITUDE |
| 19 | NOSE_VELOCITY_DIRECTION |
| 20 | NOSE_VELOCITY_MAGNITUDE |
| 21 | NOSE-LEFT_EAR |
| 22 | NOSE-RIGHT_EAR |
| 23 | NOSE-BASE_NECK |
| 24 | NOSE-LEFT_FRONT_PAW |
| 25 | NOSE-RIGHT_FRONT_PAW |
| 26 | NOSE-CENTER_SPINE |
| 27 | NOSE-LEFT_REAR_PAW |
| 28 | NOSE-RIGHT_REAR_PAW |
| 29 | NOSE-BASE_TAIL |
| 30 | NOSE-MID_TAIL |
| 31 | NOSE-TIP_TAIL |
| 32 | LEFT_EAR-RIGHT_EAR |
| 33 | LEFT_EAR-BASE_NECK |
| 34 | LEFT_EAR-LEFT_FRONT_PAW |
| 35 | LEFT_EAR-RIGHT_FRONT_PAW |
| 36 | LEFT_EAR-CENTER_SPINE |
| 37 | LEFT_EAR-LEFT_REAR_PAW |
| 38 | LEFT_EAR-RIGHT_REAR_PAW |
| 39 | LEFT_EAR-BASE_TAIL |
| 40 | LEFT_EAR-MID_TAIL |
| 41 | LEFT_EAR-TIP_TAIL |
| 42 | RIGHT_EAR-BASE_NECK |
| 43 | RIGHT_EAR-LEFT_FRONT_PAW |
| 44 | RIGHT_EAR-RIGHT_FRONT_PAW |
| 45 | RIGHT_EAR-CENTER_SPINE |
| 46 | RIGHT_EAR-LEFT_REAR_PAW |
| 47 | RIGHT_EAR-RIGHT_REAR_PAW |
| 48 | RIGHT_EAR-BASE_TAIL |
| 49 | RIGHT_EAR-MID_TAIL |
| 50 | RIGHT_EAR-TIP_TAIL |
| 51 | BASE_NECK-LEFT_FRONT_PAW |
| 52 | BASE_NECK-RIGHT_FRONT_PAW |
| 53 | BASE_NECK-CENTER_SPINE |
| 54 | BASE_NECK-LEFT_REAR_PAW |
| 55 | BASE_NECK-RIGHT_REAR_PAW |
| 56 | BASE_NECK-BASE_TAIL |
| 57 | BASE_NECK-MID_TAIL |
| 58 | BASE_NECK-TIP_TAIL |
| 59 | LEFT_FRONT_PAW-RIGHT_FRONT_PAW |
| 60 | LEFT_FRONT_PAW-CENTER_SPINE |
| 61 | LEFT_FRONT_PAW-LEFT_REAR_PAW |
| 62 | LEFT_FRONT_PAW-RIGHT_REAR_PAW |
| 63 | LEFT_FRONT_PAW-BASE_TAIL |
| 64 | LEFT_FRONT_PAW-MID_TAIL |
| 65 | LEFT_FRONT_PAW-TIP_TAIL |
| 66 | RIGHT_FRONT_PAW-CENTER_SPINE |
| 67 | RIGHT_FRONT_PAW-LEFT_REAR_PAW |
| 68 | RIGHT_FRONT_PAW-RIGHT_REAR_PAW |
| 69 | RIGHT_FRONT_PAW-BASE_TAIL |
| 70 | RIGHT_FRONT_PAW-MID_TAIL |
| 71 | RIGHT_FRONT_PAW-TIP_TAIL |
| 72 | CENTER_SPINE-LEFT_REAR_PAW |
| 73 | CENTER_SPINE-RIGHT_REAR_PAW |
| 74 | CENTER_SPINE-BASE_TAIL |
| 75 | CENTER_SPINE-MID_TAIL |
| 76 | CENTER_SPINE-TIP_TAIL |
| 77 | LEFT_REAR_PAW-RIGHT_REAR_PAW |
| 78 | LEFT_REAR_PAW-BASE_TAIL |
| 79 | LEFT_REAR_PAW-MID_TAIL |
| 80 | LEFT_REAR_PAW-TIP_TAIL |
| 81 | RIGHT_REAR_PAW-BASE_TAIL |
| 82 | RIGHT_REAR_PAW-MID_TAIL |
| 83 | RIGHT_REAR_PAW-TIP_TAIL |
| 84 | BASE_TAIL-MID_TAIL |
| 85 | BASE_TAIL-TIP_TAIL |
| 86 | MID_TAIL-TIP_TAIL |
| 87 | NOSE POINT MASK |
| 88 | LEFT_EAR POINT MASK |
| 89 | RIGHT_EAR POINT MASK |
| 90 | BASE_NECK POINT MASK |
| 91 | LEFT_FRONT_PAW POINT MASK |
| 92 | RIGHT_FRONT_PAW POINT MASK |
| 93 | CENTER_SPINE POINT MASK |
| 94 | LEFT_REAR_PAW POINT MASK |
| 95 | RIGHT_REAR_PAW POINT MASK |
| 96 | BASE_TAIL POINT MASK |
| 97 | MID_TAIL POINT MASK |
| 98 | TIP_TAIL POINT MASK |
| 99 | NOSE SPEED |
| 100 | LEFT_EAR SPEED |
| 101 | RIGHT_EAR SPEED |
| 102 | BASE_NECK SPEED |
| 103 | LEFT_FRONT_PAW SPEED |
| 104 | RIGHT_FRONT_PAW SPEED |
| 105 | CENTER_SPINE SPEED |
| 106 | LEFT_REAR_PAW SPEED |
| 107 | RIGHT_REAR_PAW SPEED |
| 108 | BASE_TAIL SPEED |
| 109 | MID_TAIL SPEED |
| 110 | TIP_TAIL SPEED |
| 111 | RIGHT FRONT PAW VELOCITY DIRECTION |
| 112 | RIGHT FRONT PAW VELOCITY MAGNITUDE |

**Figure S3:**
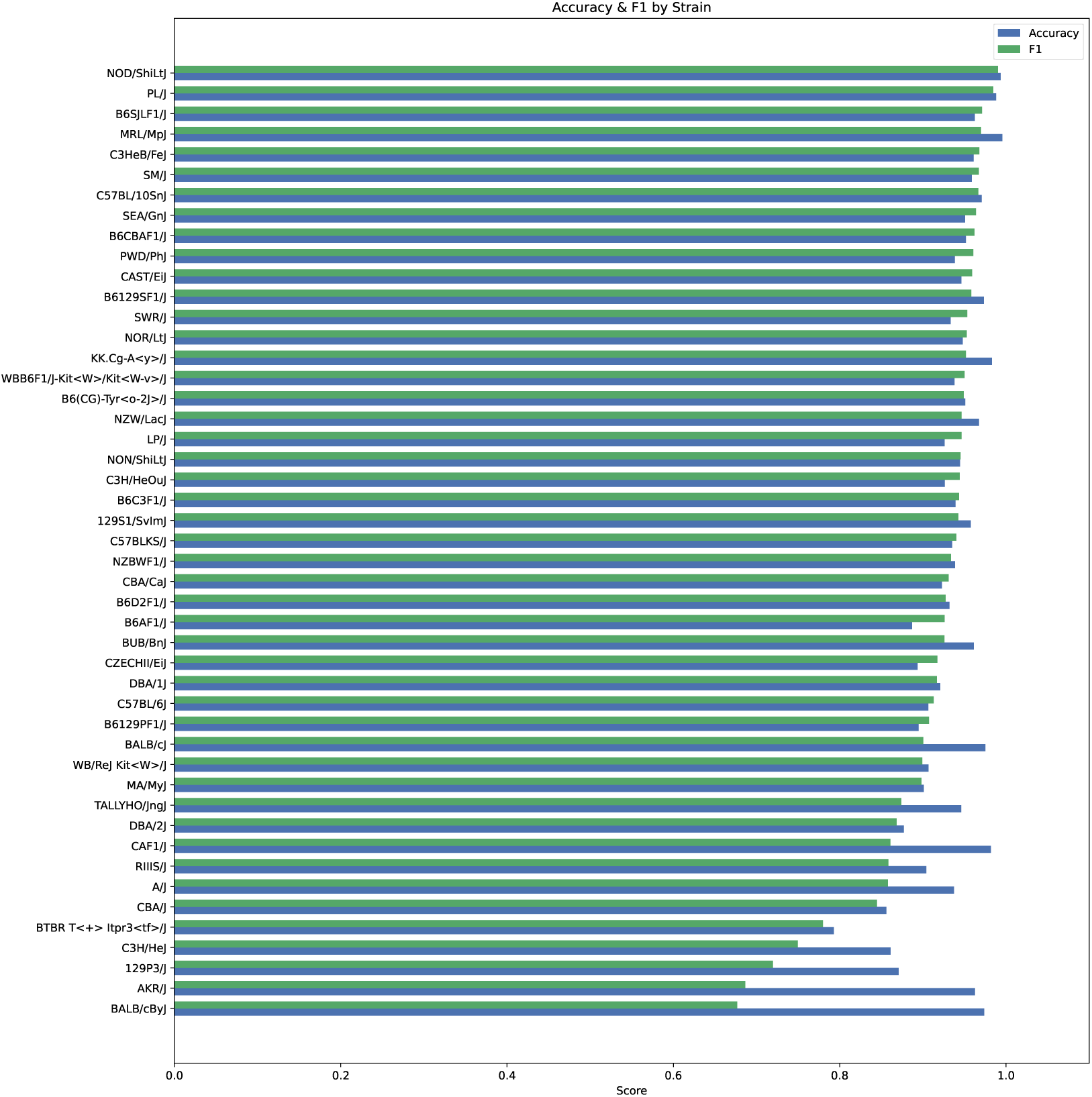
Per strain F1 score and accuracy for Grooming dataset

**Figure S4:**
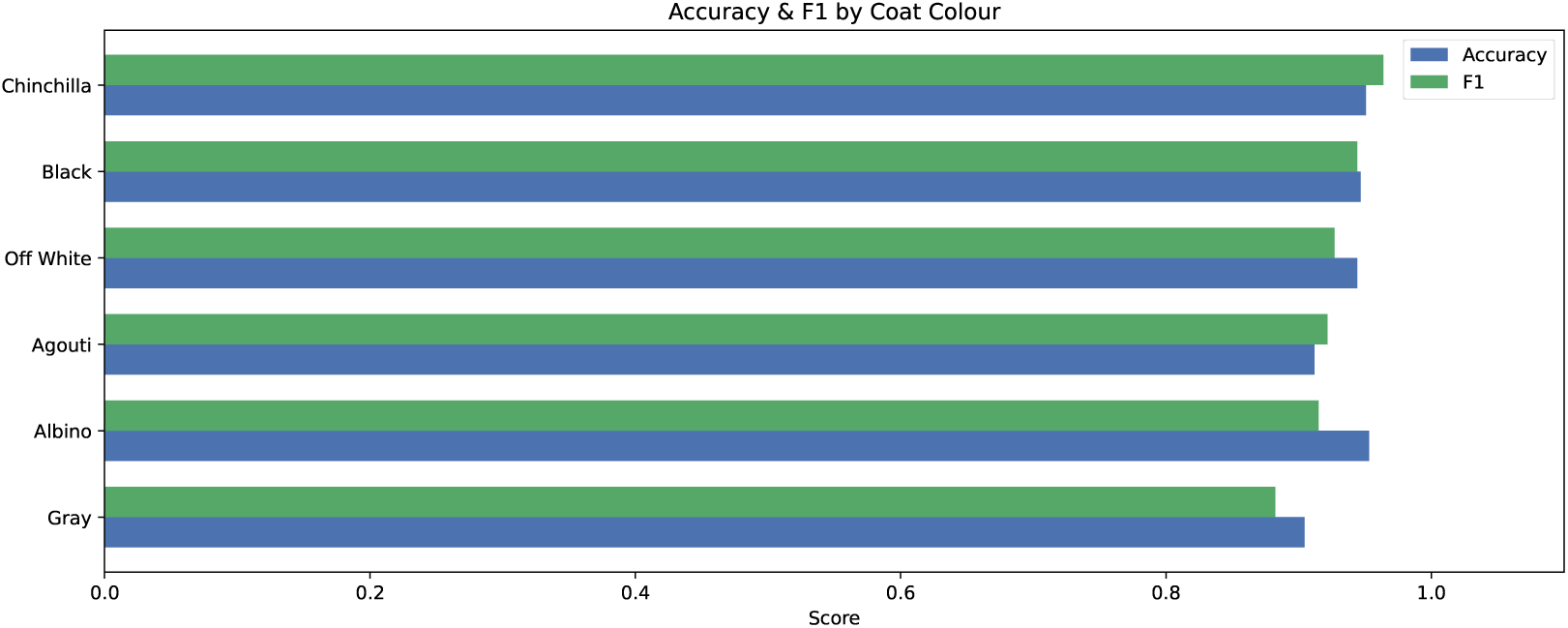
F1 score and accuracy for mice with different coat colors in the grooming dataset

**Table S3:** Behavioral phenotypes annotated by different annotators (A1, A2). Each phenotype measures a specific metric related to bouts of the indicated behavior during the first 55 minutes of the video averaged across all the analyzed videos.

| Phenotype | Description |
| --- | --- |
| A2_TL_T55_avgLen | Average bout length predicted by left turn behavior classifier (trained on labels by annotator 2) during first 55 minutes. |
| A2_TL_T55_nBouts | Number of left turn bouts predicted by left turn behavior classifier (trained on labels by annotator 2) during first 55 minutes. |
| A2_TL_T55_duration | Total duration predicted by left turn behavior classifier (trained on labels by annotator 2) during first 55 minutes. |
| A1_TR_T55_avgLen | Average bout length predicted by right turn behavior classifier (trained on labels by annotator 1) during first 55 minutes. |
| A1_TR_T55_nBouts | Number of right turn bouts predicted by left turn behavior classifier (trained on labels by annotator 1) during first 55 minutes. |
| A1_TR_T55_duration | Total duration predicted by right turn behavior classifier (trained on labels by annotator 1) during first 55 minutes. |
| A1_TL_T55_avgLen | Average bout length predicted by left turn behavior classifier (trained on labels by annotator 1) during first 55 minutes. |
| A1_TL_T55_nBouts | Number of left turn bouts predicted by left turn behavior classifier (trained on labels by annotator 1) during first 55 minutes. |
| A1_TL_T55_duration | Total duration predicted by left turn behavior classifier (trained on labels by annotator 1) during first 55 minutes. |
| SR_T55_avgLen | Average bout length predicted by scratch behavior classifier during first 55 minutes. |
| SR_T55_nBouts | Number of bouts predicted by scratch behavior classifier during first 55 minutes. |
| SR_T55_duration | Total duration predicted by scratch behavior classifier during first 55 minutes. |
| RUS_T55_avgLen | Average bout length predicted by rearing unsupported behavior classifier during first 55 minutes. |
| RUS_T55_nBouts | Number of bouts predicted by rearing unsupported behavior classifier during first 55 minutes. |
| RUS_T55_duration | Total duration predicted by rearing unsupported behavior classifier during first 55 minutes. |
| RS_T55_avgLen | Average bout length predicted by rearing supported behavior classifier during first 55 minutes. |
| RS_T55_nBouts | Number of bouts predicted by rearing supported behavior classifier during first 55 minutes. |
| RS_T55_duration | Total duration predicted by rearing supported behavior classifier during first 55 minutes. |
| GR_T55_avgLen | Average bout length predicted by grooming behavior classifier during first 55 minutes. |
| GR_T55_nBouts | Number of bouts predicted by grooming behavior classifier during first 55 minutes. |
| GR_T55_duration | Total duration predicted by grooming behavior classifier during first 55 minutes. |

**Figure S5:**
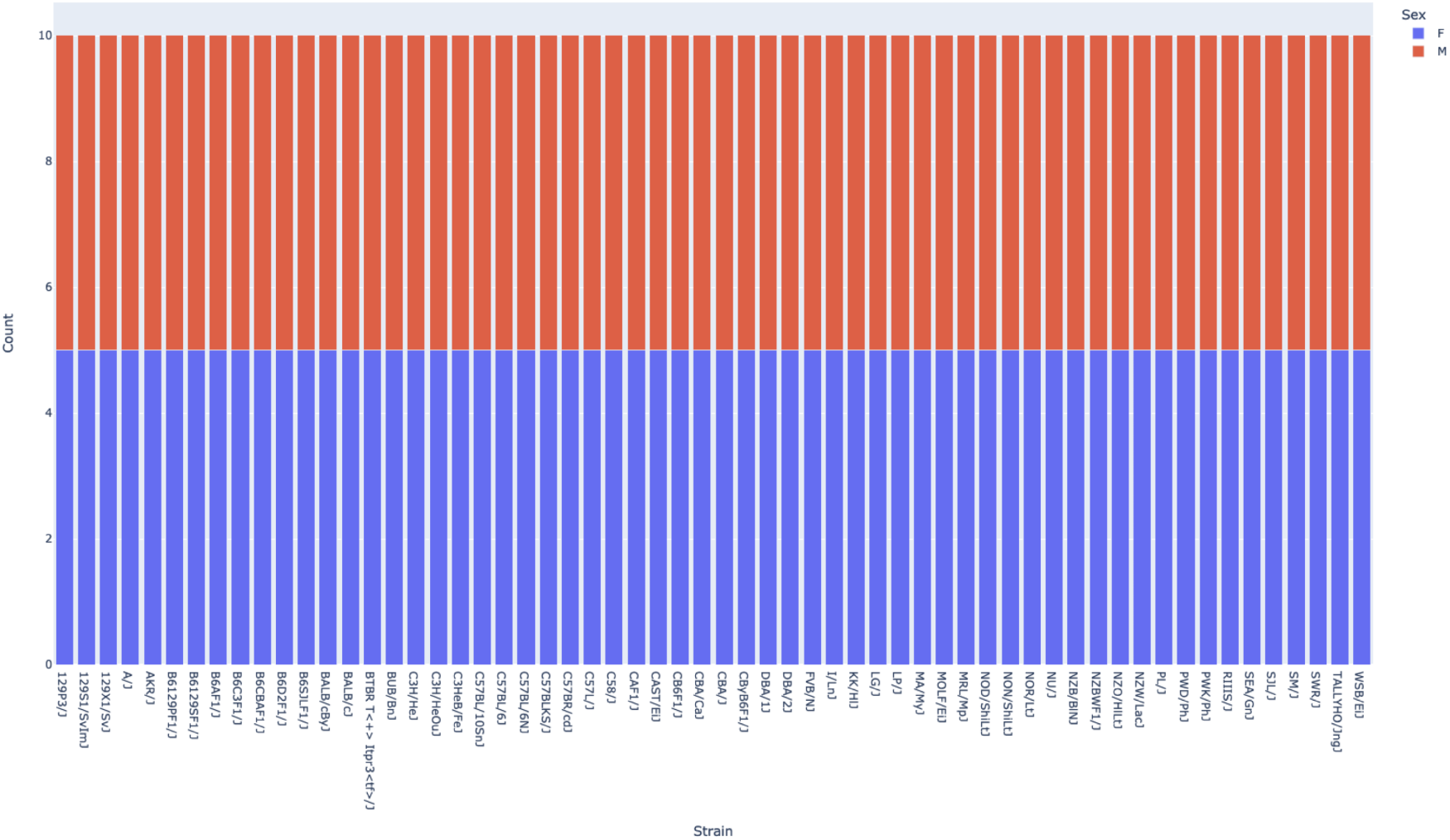
JABS 600 Strain Distribution by Sex

**Figure S6:**
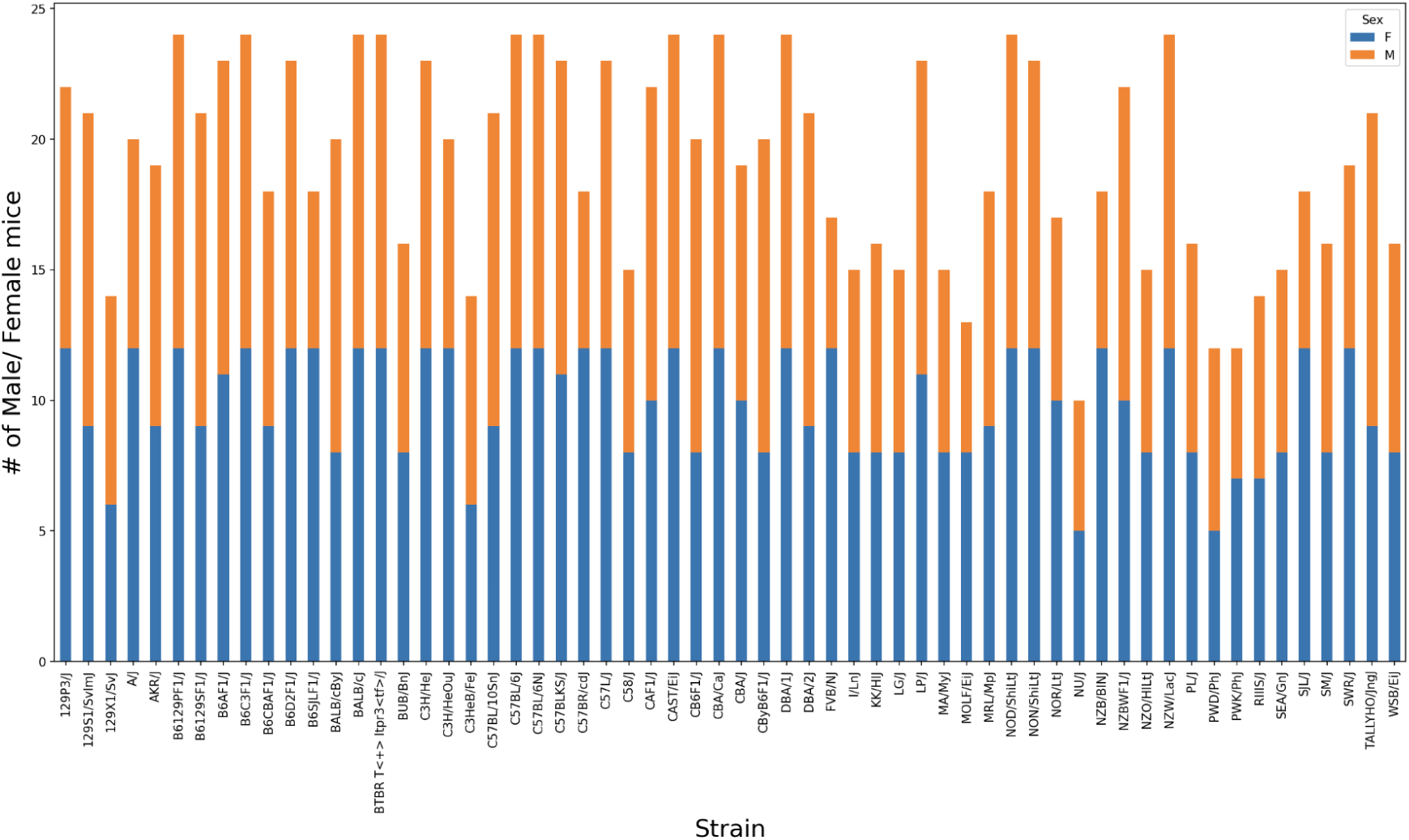
JABS 1200 Strain Distribution by Sex

**Table S5:** Classifiers trained by JABS with their respective window sizes and F1 scores.

| Classifier | Window Size | Accuracy | F1 score | Precision | Recall |
| --- | --- | --- | --- | --- | --- |
| Escape | 2 | 0.96 | 0.87 | 0.86 | 0.93 |
| Grooming | 60 | 0.94 | 0.92 | 0.92 | 0.93 |
| Rearing supported | 5 | 0.95 | 0.93 | 0.95 | 0.92 |
| Rearing unsupported | 10 | 0.87 | 0.78 | 0.88 | 0.77 |
| Turn Left (Annotator-1) | 5 | 0.91 | 0.78 | 0.84 | 0.76 |
| Turn Left (Annotator-2) | 5 | 0.91 | 0.89 | 0.87 | 0.93 |
| Turn Right (Annotator-1) | 20 | 0.92 | 0.85 | 0.85 | 0.88 |
| Turn Right (Annotator-2) | 5 | 0.90 | 0.87 | 0.85 | 0.90 |

**Figure S7:**
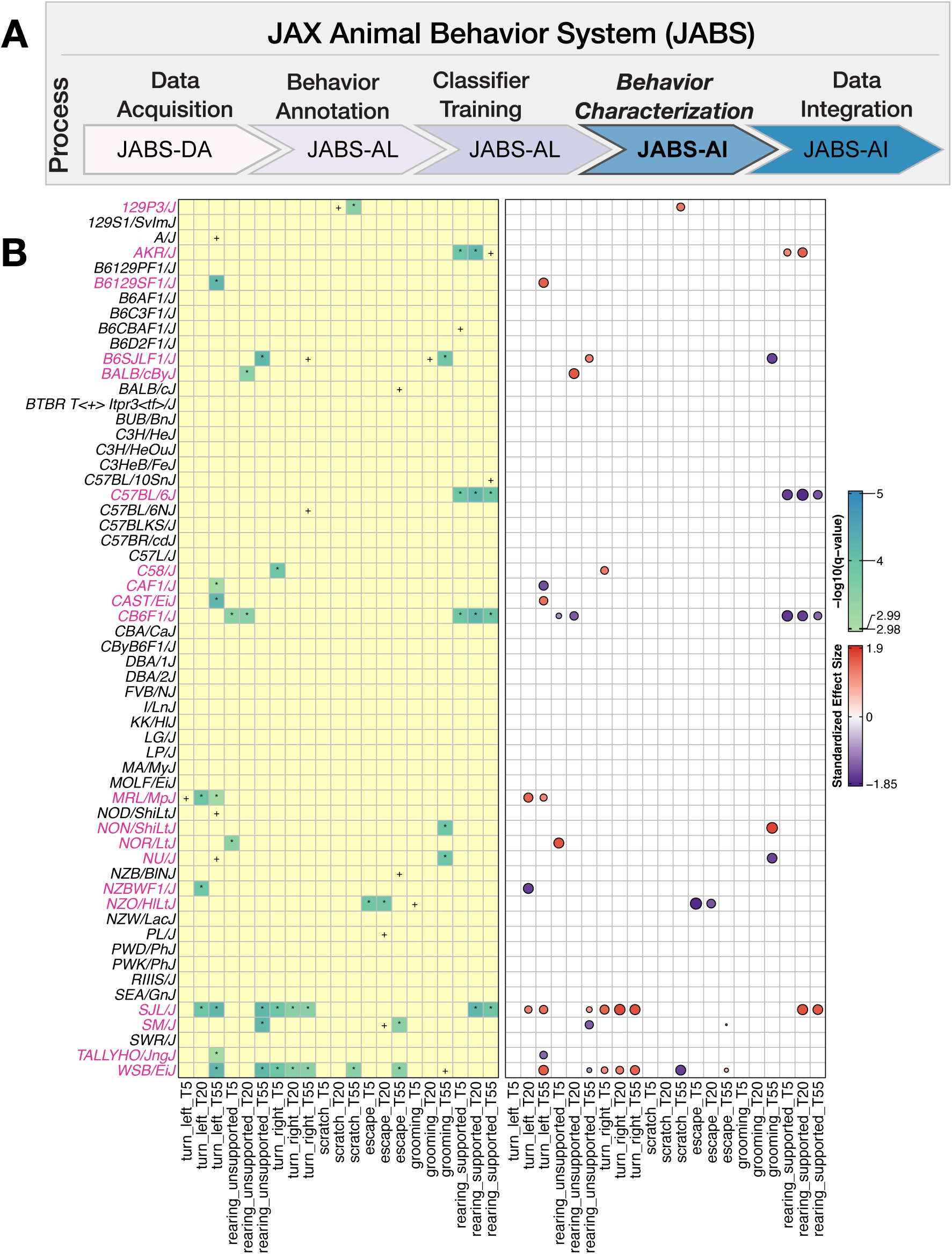
JABS behavior characterization module: Univariate analysis captures the combined effect of sex and strain on the aggregate phenotypes using JABS600 dataset: (A) JABS pipeline highlighting individual steps towards automated behavioral quantification. (B) The LOD scores (−*log*_10_(*q_value_*)) and effect sizes are shown at left and right panels, respectively. In the left panel, the number of *s represents the strength of evidence against the null hypothesis of no sex effect, while + represents a suggestive effect. In the right panel, the color (red for female and blue for male) and area of the circle (area being proportional to the size of the effect) represent the direction and magnitude of the effect size. Strains with a sex difference in at least one of the aggregated phenotypes are colored pink.

## References

1. Gomez-Marin, A., Paton, J. J., Kampff, A. R., Costa, R. M. & Mainen, Z. F. Big behavioral data: psychology, ethology and the foundations of neuroscience. Nature neuroscience 17, 1455–1462 (2014).

2. Datta, S. R., Anderson, D. J., Branson, K., Perona, P. & Leifer, A. Computational neuroethology: a call to action. Neuron 104, 11–24 (2019).

3. Pereira, T. D., Shaevitz, J. W. & Murthy, M. Quantifying behavior to understand the brain. Nature neuroscience, 1–13 (2020).

4. Luxem, K. et al. Open-source tools for behavioral video analysis: Setup, methods, and best practices. Elife 12, e79305 (2023).

5. Anderson, D. J. & Perona, P. Toward a science of computational ethology. Neuron 84, 18–31 (2014).

6. Mathis, A., Schneider, S., Lauer, J. & Mathis, M. W. A primer on motion capture with deep learning: principles, pitfalls, and perspectives. Neuron 108, 44–65 (2020).

7. Choi, J. D. & Kumar, V. A new era in quantification of animal social behaviors. en. Neurosci. Biobehav. Rev. 157, 105528 (Feb. 2024).

8. Geuther, B. Q. et al. Robust mouse tracking in complex environments using neural networks. Communications biology 2, 124 (2019).

9. Geuther, B. Q. et al. Action detection using a neural network elucidates the genetics of mouse grooming behavior. Elife 10, e63207 (2021).

10. Sheppard, K. et al. Stride-level analysis of mouse open field behavior using deep-learning-based pose estimation. Cell Reports 38, 110231 (2022).

11. Hession, L. E., Sabnis, G., Churchill, G. A. & Kumar, V. A machine vision based frailty index for mice. bioRxiv (2021).

12. Guzman, M., Geuther, B. Q., Sabnis, G. S. & Kumar, V. Highly accurate and precise determination of mouse mass using computer vision. Patterns (2023).

13. Sabnis, G. S., Hession, L. E., Kim, K., Beierle, J. A. & Kumar, V. A high-throughput machine vision-based univariate scale for pain and analgesia in mice. bioRxiv, 2022–12 (2022).

14. Hession, L. E., Sabnis, G. S., Churchill, G. A. & Kumar, V. A machine-vision-based frailty index for mice. Nature aging 2, 756–766 (2022).

15. Sabnis, G., et al. Visual detection of seizures in mice using supervised machine learning. bioRxiv (2024).

16. Isik, S. & Unal, G. Open-source software for automated rodent behavioral analysis. Frontiers in Neuroscience 17, 1149027 (2023).

17. Mathis, A. et al. DeepLabCut: markerless pose estimation of user-defined body parts with deep learning. Nature neuroscience 21, 1281 (2018).

18. Pereira, T. D. et al. SLEAP: A deep learning system for multi-animal pose tracking. Nature methods 19, 486–495 (2022).

19. Goodwin, N. L. et al. Simple Behavioral Analysis (SimBA) as a platform for explainable machine learning in behavioral neuroscience. Nature Neuroscience 27, 1411–1424 (2024).

20. Segalin, C. et al. The Mouse Action Recognition System (MARS) software pipeline for automated analysis of social behaviors in mice. Elife 10 (2021).

21. Kabra, M., Robie, A. A., Rivera-Alba, M., Branson, S. & Branson, K. JAABA: interactive machine learning for automatic annotation of animal behavior. Nature methods 10, 64 (2013).

22. Hsu, A. I. & Yttri, E. A. B-SOiD, an open-source unsupervised algorithm for identification and fast prediction of behaviors. Nature communications 12, 5188 (2021).

23. Bohnslav, J. P. et al. DeepEthogram, a machine learning pipeline for supervised behavior classification from raw pixels. elife 10, e63377 (2021).

24. Kumar, V. et al. Second-generation high-throughput forward genetic screen in mice to isolate subtle behavioral mutants. Proceedings of the National Academy of Sciences 108, 15557–15564. ISSN: 0027-8424 (2011).

25. Council, N. R., et al. Guide for the care and use of laboratory animals (2011).

26. Myatt, T. A. et al. A study of indoor carbon dioxide levels and sick leave among office workers. Environmental Health 1, 1–10 (2002).

27. Mexas, A. M., Brice, A. K., Caro, A. C., Hillanbrand, T. S. & Gaertner, D. J. Nasal histopathology and intracage ammonia levels in female groups and breeding mice housed in static isolation cages. Journal of the American Association for Laboratory Animal Science 54, 478–486 (2015).

28. Fawcett, A. & Rose, M. Guidelines for the housing of mice in scientific institutions. *Animal Welfare Unit, NSW Department of Primary Industries*, West Pennant Hills. Anim Res Rev Panel 1, 1–43 (2012).

29. Gamble, M. & Clough, G. Ammonia build-up in animal boxes and its effect on rat tracheal epithelium. Laboratory Animals 10, 93–104 (1976).

30. Ferrecchia, C. E., Jensen, K. & Van Andel, R. Intracage ammonia levels in static and individually ventilated cages housing C57BL/6 mice on 4 bedding substrates. Journal of the American Association for Laboratory Animal Science 53, 146–151 (2014).

31. Tepper, J. S., Weiss, B. & Wood, R. W. Alterations in behavior produced by inhaled ozone or ammonia. Fundamental and Applied Toxicology 5, 1110–1118 (1985).

32. Easterly, M. E., Foltz, J. & Paulus, M. J. Body condition scoring: comparing newly trained scorers and micro-computed tomography imaging. LAB ANIMAL-NEW YORK*-* 30, 46–49 (2001).

33. Hickman, D. L. & Swan, M. Use of a body condition score technique to assess health status in a rat model of polycystic kidney disease. Journal of the American Association for Laboratory Animal Science 49, 155–159 (2010).

34. Lovasz, R. M., Marks, D. L., Chan, B. K. & Saunders, K. E. Effects on Mouse Food Consumption After Exposure to Bedding from Sick Mice or Healthy Mice. Journal of the American Association for Laboratory Animal Science 59, 687–694 (2020).

35. Green, E. L. Biology of the laboratory mouse (1966).

36. Biderman, D. et al. Lightning Pose: improved animal pose estimation via semi-supervised learning, Bayesian ensembling and cloud-native open-source tools. Nature methods 21, 1316–1328 (2024).

37. Chen, T. & Guestrin, C. *Xgboost: A scalable tree boosting system* in *Proceedings of the 22nd acm sigkdd international conference on knowledge discovery and data mining* (2016), 785–794.

38. Ho, T. K. *Random decision forests* in Proceedings of 3rd international conference on document analysis and recognition 1 (1995), 278–282.

39. Breiman, L. Random forests. Machine learning 45, 5–32 (2001).

40. Friedman, J. H. Greedy function approximation: a gradient boosting machine. Annals of statistics, 1189–1232 (2001).

41. Mathis, A. et al. DeepLabCut: markerless pose estimation of user-defined body parts with deep learning. Nature Neuroscience 21 (Sept. 2018).

42. Kaufman, A. B. & Rosenthal, R. Can you believe my eyes? The importance of interobserver reliability statistics in observations of animal behaviour. Animal Behaviour 78, 1487–1491 (2009).

43. Tjandrasuwita, M., Sun, J. J., Kennedy, A., Chaudhuri, S. & Yue, Y. Interpreting expert annotation differences in animal behavior. arXiv preprint arXiv:2106.06114 (2021).

44. McHugh, M. L. Interrater reliability: the kappa statistic. Biochemia medica 22, 276–282 (2012).

45. Feichtenhofer, C., Fan, H., Malik, J. & He, K. Slowfast networks for video recognition in Proceedings of the IEEE International Conference on Computer Vision (2019), 6202–6211.

46. Kalogeiton, V., Weinzaepfel, P., Ferrari, V. & Schmid, C. Action tubelet detector for spatiotemporal action localization in Proceedings of the IEEE International Conference on Computer Vision (2017), 4405–4413.

47. Ryan, B. C., Young, N. B., Crawley, J. N., Bodfish, J. W. & Moy, S. S. Social deficits, stereotypy and early emergence of repetitive behavior in the C58/J inbred mouse strain. Behavioural Brain Research 208, 178–188. ISSN: 0166-4328. https://www.sciencedirect.com/science/article/pii/S0166432809007086 (2010).

48. Blick, M. G. et al. Novel object exploration in the C58/J mouse model of autistic-like behavior. Behavioural Brain Research 282, 54–60. ISSN: 0166-4328. https://www.sciencedirect.com/science/article/pii/S0166432814008249 (2015).

49. Ye, S. et al. SuperAnimal pretrained pose estimation models for behavioral analysis. Nature communications 15, 5165 (2024).

50. Pendergrass, S. et al. The use of phenome-wide association studies (PheWAS) for exploration of novel genotype-phenotype relationships and pleiotropy discovery. Genetic epidemiology 35, 410–422 (2011).

51. Weinreb, C. et al. Keypoint-MoSeq: parsing behavior by linking point tracking to pose dynamics. Nature Methods 21, 1329–1339 (2024).

52. Luxem, K. et al. Identifying behavioral structure from deep variational embeddings of animal motion. Communications Biology 5, 1267 (2022).

53. Berman, G. J., Choi, D. M., Bialek, W. & Shaevitz, J. W. Mapping the stereotyped behaviour of freely moving fruit flies. Journal of The Royal Society Interface 11, 20140672 (2014).

54. Blau, A. et al. A study of animal action segmentation algorithms across supervised, unsupervised, and semi-supervised learning paradigms. ArXiv, arXiv–2407 (2024).

55. Rosenberg, M., Zhang, T., Perona, P. & Meister, M. Mice in a labyrinth: Rapid learning, sudden insight, and efficient exploration. DOI:10.1101/2021.01.14.426746v1 (2021).

56. Arakawa, H., Blanchard, D. C. & Blanchard, R. J. Colony formation of C57BL/6J mice in visible burrow system: identification of eusocial behaviors in a background strain for genetic animal models of autism. Behavioural brain research 176, 27–39 (2007).

57. Zhou, X. & Stephens, M. Genome-wide efficient mixed-model analysis for association studies. Nature genetics 44, 821–824 (2012).

